# The widely used *Ucp1-Cre^Evdr^* transgene elicits complex developmental and metabolic phenotypes

**DOI:** 10.1101/2023.10.20.563165

**Authors:** Manasi Suchit Halurkar, Oto Inoue, Rajib Mukherjee, Christian Louis Bonatto Paese, Molly Duszynski, Samantha A. Brugmann, Hee-Woong Lim, Joan Sanchez-Gurmaches

**Author notes:** Allen Institute for Brain Science, Seattle, WA 98109, USA.

## Abstract

Bacterial artificial chromosome transgenic models, including most *Cre-recombinases*, enable potent interrogation of gene function *in vivo* but require rigorous validation as limitations emerge. Due to its high relevance to metabolic studies, we performed comprehensive analysis of the *Ucp1-Cre^Evdr^* line which is widely used for brown fat research. Hemizygotes exhibited major brown and white fat transcriptomic dysregulation, indicating potential altered tissue function. *Ucp1-Cre^Evdr^*homozygotes also show high mortality, growth defects, and craniofacial abnormalities. Mapping the transgene insertion site revealed insertion in chromosome 1 accompanied by large genomic alterations disrupting several genes expressed in a range of tissues. Notably, *Ucp1-Cre^Evdr^* transgene retains an extra *Ucp1* gene copy that may be highly expressed under high thermogenic burden. Our multi-faceted analysis highlights a complex phenotype arising from the presence of the *Ucp1-Cre^Evdr^* transgene independently of the intended genetic manipulations. Overall, comprehensive validation of transgenic mice is imperative to maximize discovery while mitigating unexpected, off-target effects.

**Highlights:**

- Hemizygous *Ucp1-Cre^Evdr^* mice exhibit substantial brown and white fat tissue dysregulation.
- Homozygous *Ucp1-Cre^Evdr^* mice display high mortality, growth defects, and craniofacial abnormalities.
- The *Ucp1-Cre^Evdr^* transgene integration resulted in major genomic disruptions affecting multiple genes.
- The *Ucp1-Cre^Evdr^* transgene retains a possibly functional extra *Ucp1* copy.

## Introduction

Mouse transgenic models such as overexpressors, reporters, and Cre-recombinases empower spatial and temporal genetic manipulation enabling unparalleled interrogation of gene function *in vivo*. These models have driven discoveries across biological scales, from molecular processes to whole body physiology and have become indispensable for elucidating the foundations of health and disease.

Most transgenic mouse alleles in use today have been generated via bacterial artificial chromosome (BAC) technology^1,2^. This involves inserting sequences of interest (e.g. Cre-recombinase) into BAC plasmids (∼150-350 kb) containing regulatory elements that confer spatiotemporal expression. For instance, insertion after a promoter sequence permits cell type-or stage-specific Cre-recombinase expression. The modified BAC, including contextual sequences, is then randomly integrated into the host genome in a nonspecific, stochastic manner, typically forming a multicopy concatemer^3^.

While revolutionary, *Cre*-driver lines generated by BAC transgenesis carry potential limitations that are rarely investigated. Validation of *Cre*-drivers is usually restricted to verification of specific expression in the targeted cell type. Initiatives like the CrePortal^4,5^ have been invaluable for collating *Cre* expression data and provide a valuable resource to guide appropriate use of *Cre*-drivers. Yet other limitations associated with BAC transgenesis are rarely examined: (1) The insertion site of transgenes are mostly unknown; only 5.03% and 3.40% of all transgenic and *Cre* alleles, respectively, have mapped integration sites collated in Mouse Genome Informatics [Figure S1A-B]; (2) insertion can result in large genomic abnormalities that are not routinely inspected^6,7,8^ and additionally the insertion may directly influence the phenotypes observed by different mechanisms^9–12^; (3) passenger sequences are virtually never reported but may lead to unintended phenotypes^13^; (4) *Cre* transgenes are largely used in hemizygosity masking phenotypes that would otherwise be evident^14,15^; (5) the common absence of *Cre*-only control groups precludes assessment of perturbations directly attributable to the presence of the *Cre* transgene or protein itself.

BAC transgenics have been instrumental for generating adipose-specific Cre driver lines to dissect the biology of the highly thermogenic brown adipose tissue (BAT) and the energy storing white adipose tissue (WAT)^16–18^. *Cre-recombinase* lines utilizing the adiponectin promoter enabled targeting of all adipocytes^19–21^. Promoter elements from *UnCoupling Protein 1* (*Ucp1*) have conferred selective *Cre-recombinase* expression in brown adipocytes. Although other BAT *Cre*-targeting tools existed^22^, and other tools to target brown adipocytes are used^23^, two *Ucp1-Cre* drivers dominate the literature currently: the constitutive *Ucp1-Cre^Evdr^* line from the Rosen lab^24^ and the tamoxifen inducible *Ucp1-CreERT2^Biat^* line from the Wolfrum lab^25^. Both lines show remarkable specificity, full penetrance, and robust activity on brown adipocytes^26–32^. Among the two, the *Ucp1-Cre^Evdr^* line has been more widely adopted, featuring in 78.85% of manuscripts in the Mouse Genome Informatics records. This skewed utilization may be due to greater accessibility in open repositories and concerns about tamoxifen effects on adipocytes^33,34^.

Despite the extensive use of the constitutive *Ucp1-Cre^Evdr^*allele, comprehensive validation of this driver line is lacking. Here, we perform an in-depth characterization of *Ucp1-Cre^Evdr^* using genetic, genomic and physiologic approaches. Transcriptomic analysis showed substantial gene expression changes in both brown and white adipose tissues of hemizygous *Ucp1-Cre^Evdr^* mice compared to wild-type littermate controls. *Ucp1-Cre^Evdr^* homozygotes show high mortality, craniofacial abnormalities, and growth retardation. Molecular characterization of the *Ucp1-Cre^Evdr^* transgene insertion site demonstrated substantial genomic alterations including disruptions of several genes at the integration locus in chromosome 1. Notably, the *Ucp1-Cre^Evdr^* transgene retains an additional *Ucp1* gene that may exhibit strong expression under high thermogenic burden. These effects suggest unintended consequence on brown adipose tissue function by *Ucp1-Cre^Evdr^*. More broadly, our study highlights the critical need for extensive validation of BAC transgenic drivers.

## Results

### *UCP1-Cre^Evdr^* Homozygosity Induces Lethality, Growth Impairment, and Craniofacial Abnormalities

While attempting to generate a *Ucp1-Cre^Evdr^* mediated deletion of a *Ucp1* floxed allele, we were unable to identify mice homozygous for the *Ucp1* floxed allele and simultaneously harboring the *Ucp1-Cre^Evdr^* transgene through standard genotyping (see below). Given that germline *Ucp1* knockout mice are viable, embryonic lethality due to *Ucp1* deficiency does not explain this bias in genotyping ratios. Thus, we ponder the question on whether the *Ucp1-Cre^Evdr^*transgene itself may underlie the observed effects.

To rigorously evaluate the *Ucp1-Cre^Evdr^* model for its use in discovery-based research, we generated control, hemizygous, and homozygous littermates by crossing *Ucp1-Cre^Evdr^* hemizygous [Figure 1A]. We designated them as controls, 1x*Ucp1-Cre^Evdr^* and 2x*Ucp1-Cre^Evdr^* mice, respectively. We reasoned that this strategy would enable comprehensive assessment of potential developmental, physiological, and molecular perturbations arising from this widely utilized *Cre* driver line.

**Figure 1:**
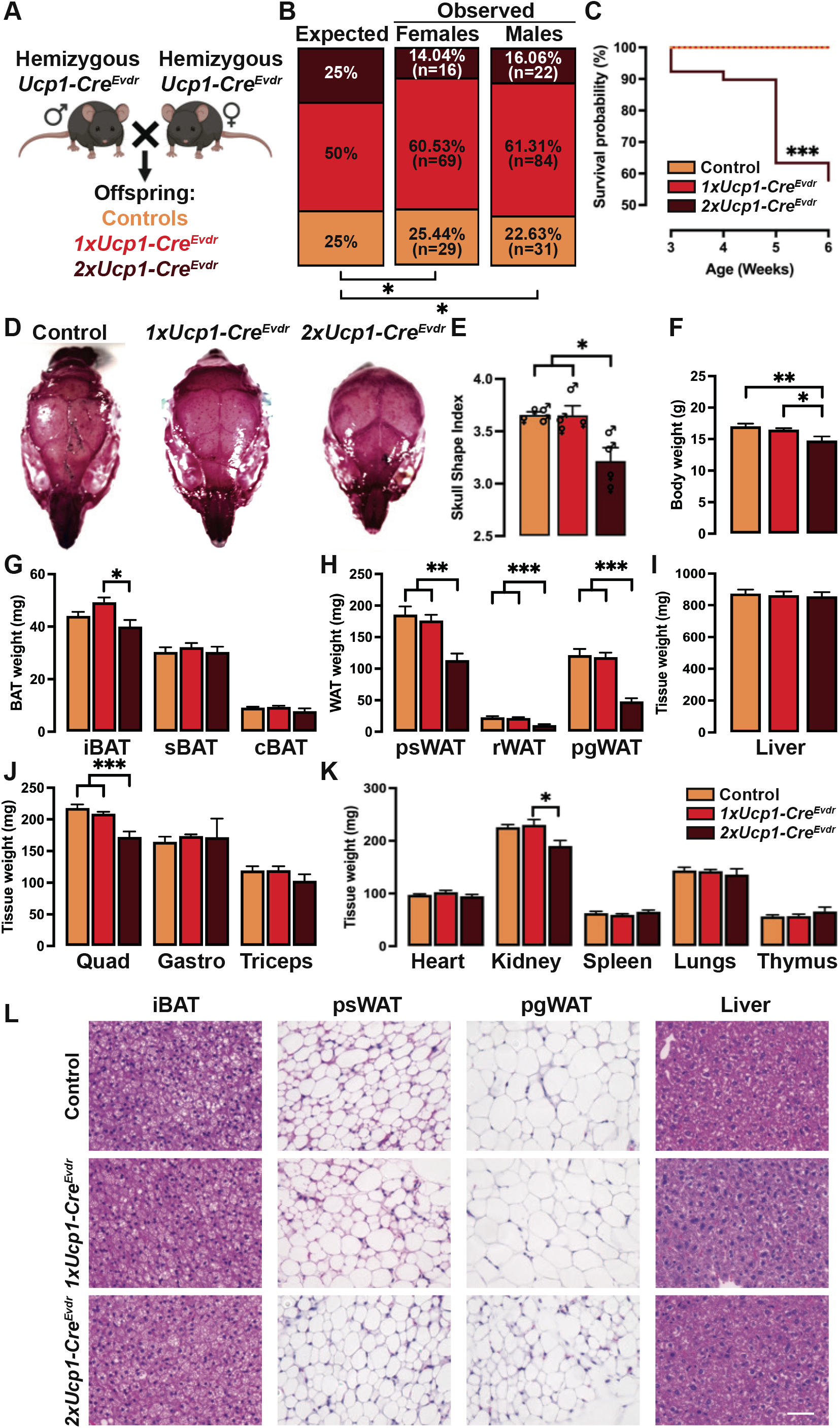
(A) Experimental strategy for the generation of control, *Ucp1-Cre^Evdr^* hemizygous (1x*Ucp1-Cre^Evdr^*) and *Ucp1-Cre^Evdr^* homozygous (2x*Ucp1-Cre^Evdr^*) mice. (B) Expected and observed offspring genotypes obtained from 1x*Ucp1-Cre^Evdr^* to 1x*Ucp1-Cre^Evdr^* crosses separated by sex. N=251 pups from 46 litters. Statistical significance was calculated using Chi-square test. (C) Kaplan-Meier survival plot of control, 1x*Ucp1-Cre^Evdr^* and 2x*Ucp1-Cre^Evdr^* mice from 3 to 6 weeks of age. n= 60 controls, 152 1x*Ucp1-Cre^Evdr^*, 23 2x*Ucp1-Cre^Evdr^*. Statistical significance was calculated using Log-rank (Mantel-Cox) test. (D) Representative photographs of alizarin red S stained skulls of control, 1x*Ucp1-Cre^Evdr^* and 2x*Ucp1-Cre^Evdr^* mice. n=2 females and 2 males. (E) Skull shape index (ration between condylobasal length and the interorbital constriction length) of control, 1x*Ucp1-Cre^Evdr^* and 2x*Ucp1-Cre^Evdr^* mice. n=2 females and 2 males. (F) Body weights of control, 1x*Ucp1-Cre^Evdr^* and 2x*Ucp1-Cre^Evdr^* females. n= 7 controls, 9 1x*Ucp1-Cre^Evdr^*, 7 2x*Ucp1-Cre^Evdr^*. (G) BAT weights of control, 1x*Ucp1-Cre^Evdr^* and 2x*Ucp1-Cre^Evdr^* females. n= 7 controls, 9 1x*Ucp1-Cre^Evdr^*, 5 2x*Ucp1-Cre^Evdr^*. (H) WAT weights of control, 1x*Ucp1-Cre^Evdr^* and 2x*Ucp1-Cre^Evdr^* females. n= 7 controls, 9 1x*Ucp1-Cre^Evdr^*, 5 2x*Ucp1-Cre^Evdr^*. (I) Liver weight of control, 1x*Ucp1-Cre^Evdr^* and 2x*Ucp1-Cre^Evdr^* females. n= 7 controls, 9 1x*Ucp1-Cre^Evdr^*, 5 2x*Ucp1-Cre^Evdr^*. (J) Skeletal muscles weight of control, 1x*Ucp1-Cre^Evdr^* and 2x*Ucp1-Cre^Evdr^* females. n= 7 controls, 9 1x*Ucp1-Cre^Evdr^*, 5 2x*Ucp1-Cre^Evdr^*. (K) Other organs weight of control, 1x*Ucp1-Cre^Evdr^* and 2x*Ucp1-Cre^Evdr^* females. n= 7 controls, 9 1x*Ucp1-Cre^Evdr^*, 5 2x*Ucp1-Cre^Evdr^*. (L) Representative H&E images of fat depots and liver from control, 1x*Ucp1-Cre^Evdr^* and 2x*Ucp1-Cre^Evdr^* females. n= 4 per genotype. Unless otherwise noted, data are mean + SEM and statistical significance was calculated using one-way ANOVA followed by Tukey’s multiple comparisons test. *P < 0.05, **P < 0.01, ***P < 0.001.

To unambiguously discriminate transgene copy number, we developed a quantitative copy number assay to detect *Cre* in genomic DNA rather than relying on endpoint PCR genotyping [Figure S1C]. At three weeks of age, we find significantly fewer 2x*Ucp1-Cre^Evdr^*mice than the expected Mendelian ratio of 25% [Figure 1B].

Specifically, only 14.04% of females and 16.06% of males are homozygous across 251 pups from 46 litters [Figure 1B]. Analysis of both sexes together reveals that 2x*Ucp1-Cre^Evdr^*comprises just 15.14% of the offspring, reflecting approximately 60% survival [Figure S1D]. Sex distribution is unaffected, indicating no differential penetrance between sexes [Figure S1E]. Moreover, over 40% of 2x*Ucp1-Cre^Evdr^* die spontaneously from 3 to 6 weeks of age, while 1x*Ucp1-Cre^Evdr^*and controls show no mortality [Figure 1C]. This mortality phenotype occurs with no indication of malaise in 2x*Ucp1-Cre^Evdr^* mice. The dramatic reduction in viability and high postnatal lethality in 2x*Ucp1-Cre^Evdr^*mice suggests profound biological perturbations by the *Ucp1-Cre^Evdr^*transgene.

To understand the potential effects of *Ucp1-Cre^Evdr^*, we next examined body weights of controls, 1x*Ucp1-Cre^Evdr^* and 2x*Ucp1-Cre^Evdr^*mice from three to six weeks old. 1x*Ucp1-Cre^Evdr^* female and male mice are indistinguishable from controls [Figure S1F-G]. However, female and male 2x*Ucp1-Cre^Evdr^* mice display 15% and 15-19%, respectively, lower body weights from 3-6 weeks of age compared to controls and hemizygotes [Figure S1F-G]. Additionally, 2x*Ucp1-Cre^Evdr^* appeared to have calvarial defects. To more carefully characterize this dysmorphology, we dissected the heads of six-week-old controls, 1x*Ucp1-Cre^Evdr^* and 2x*Ucp1-Cre^Evdr^* mice and performed Alizarin Red staining. As suspected, 2x*Ucp1-Cre^Evdr^* mice have a more domed, less elongated skull [Figure 1D]. Specifically, the frontal bones appear reduced, while the parietal bones appear increased in size. This dysmorphology resulted in a significantly reduced condylobasal to interorbital constriction length in 2x*Ucp1-Cre^Evdr^* mice [Figure 1E]. Since frontal bones are neural crest derived and parietal bones are mesodermally-derived, this may indicate a differential effect of the homozygosity of the *Ucp1-Cre^Evdr^* transgene in the development or cross-communication of these two populations^35,36^. Together, these data unveil craniofacial dysmorphologies and growth retardation in 2x*Ucp1-Cre^Evdr^* mice.

To better understand the effects of the *Ucp1-Cre^Evdr^* transgene on mouse growth, we performed comprehensive tissue dissections at 6 weeks. Despite lower total body mass in 2x*Ucp1-Cre^Evdr^* females [Figure 1F], dissection of individual fat and lean tissues show that body weight reduction is surprisingly not due to a homogeneously global reduction of weight of each independent tissue. In particular, 2x*Ucp1-Cre^Evdr^*females show no change in BAT depots weights, including interscapular (iBAT), subscapular (sBAT) and cervical (cBAT) compared to control littermates [Figure 1G]. However, posterior subcutaneous or inguinal (psWAT), retroperitoneal (rWAT) and perigonadal (pgWAT) WAT depots are severely impacted, with 39%, 53% and 60% decrease in weight respectively [Figure 1H]. Beyond WAT, only quadriceps mass differs in 2x*Ucp1-Cre^Evdr^* females compared to controls [Figure 1I-K]. Male homozygotes also exhibit similar decrease in body weight and dramatic WAT depletion along with reductions in liver, quadriceps and gastrocnemius mass [Figure S1H-M]. Thus, tissue-specific effects underlie the global growth retardation in 2x*Ucp1-Cre^Evdr^*.

Histological analysis of iBAT and psWAT reveals no major changes between genotypes in either females or males [Figure 1L, S1P]. However, adipocytes in pgWAT of 2x*Ucp1-Cre^Evdr^* females and males appear to be smaller in size [Figure 1L, S1P]. This suggests that the changes in psWAT and pgWAT weights may be due to different mechanisms involving the generation of adipocytes or control of their size. We next investigated whether aberrant *Cre* expression could explain the dramatic WAT defects in 2x*Ucp1-Cre^Evdr^*. As expected, *Cre* mRNA is undetectable in all fat depots of control mice [Figure S1N-O]. In female iBAT, *Cre* expression correlates with transgene copy number, with 3.16-fold higher levels in 2x*Ucp1-Cre^Evdr^* than 1x*Ucp1-Cre^Evdr^*. However, in psWAT and pgWAT of 2x*Ucp1-Cre^Evdr^* females, *Cre* levels remain hardly detectable and unchanged compared to 1x*Ucp1-Cre^Evdr^* [Figure S1N]. Analysis of male fat depots show similar results [Figure S1O]. This tissue distribution expression argues against *Cre* misexpression driving WAT perturbations in 2x*Ucp1-Cre^Evdr^* mice.

### The *UCP1-Cre^Evdr^* Transgene is Inserted in Chromosome 1, Disrupts Genomic Integrity, and Harbors an Extra *Ucp1* Gene Copy

To date, the genomic integration site and structure of the *Ucp1-Cre^Evdr^* transgene are unknown. To elucidate the integration site of the *Ucp1-Cre^Evdr^* transgene, we performed targeted locus amplification (TLA, Cergentis) in a hemizygous male. TLA is an unbiased, genome-wide method that utilizes sequence-specific inverse PCR of a circularized genomic DNA library following *NlaIII* fragmentation and crosslinking. Subsequent deep sequencing of PCR products enables mapping of the transgene insertion locus^37^.

TLA using Cre-specific primers reveals the *Ucp1-Cre^Evdr^* transgene integrated into chromosome 1, cytoband A5 [Figure S2A, 2A]. As expected, *Cre* primers also detects homology near the endogenous *Ucp1* locus in chromosome 8, indicating inclusion of surrounding *Ucp1* genomic sequences in the transgene [Figure 2A]. Primer pairs surrounding *Ucp1* produces high signal levels at the *Ucp1* locus [Figure S2A-B]. TLA maps the concatemer insertion site of *Ucp1-Cre^Evdr^* to chr1:20,962,125-21,016,858 [Figure 2B]. Integration induces a ∼54 kb deletion flanking the insertion sites, along with a 3’ ∼280 kb inversion [Figure 2B]. This directly deletes or inverts the entirety or large portions of 4 genes (*Paqr8*, *Efhc1*, *Tram2*, *Tmem14a*) [Figure 2B]. Additionally, the concatemer localizes in close proximity to other 7 genes (*Il17a*, *Il17f*, *Mcm3*, *Gsta3*, *Khdc1a*, *Khdc1c*, *Khdc1b*). Several noncoding sequences within or in close proximity to the concatemer may also be affected [Figure 2B]. The majority of the genes directly affected or in close proximity to the *Ucp1-Cre^Evdr^* concatemer exhibit low expression in adipose tissues, but they are selectively highly expressed in an array of other tissues [Figure 2C]. Knockout mouse models have not been generated for each coding gene affected by the *Ucp1-Cre^Evdr^* transgene [Figure 2D]. However, out those generated, only the *Mcm3* knockout mice show prenatal lethality with complete penetrance^38^ [Figure 2D]. Although this is quite distinct from what we observe in 2x*Ucp1-Cre^Evdr^*mice [Figure 1B-C, S1D-E], the genomic disruption induced by the *Ucp1-Cre^Evdr^*concatemer may contribute to the above identified perturbations.

**Figure 2:**
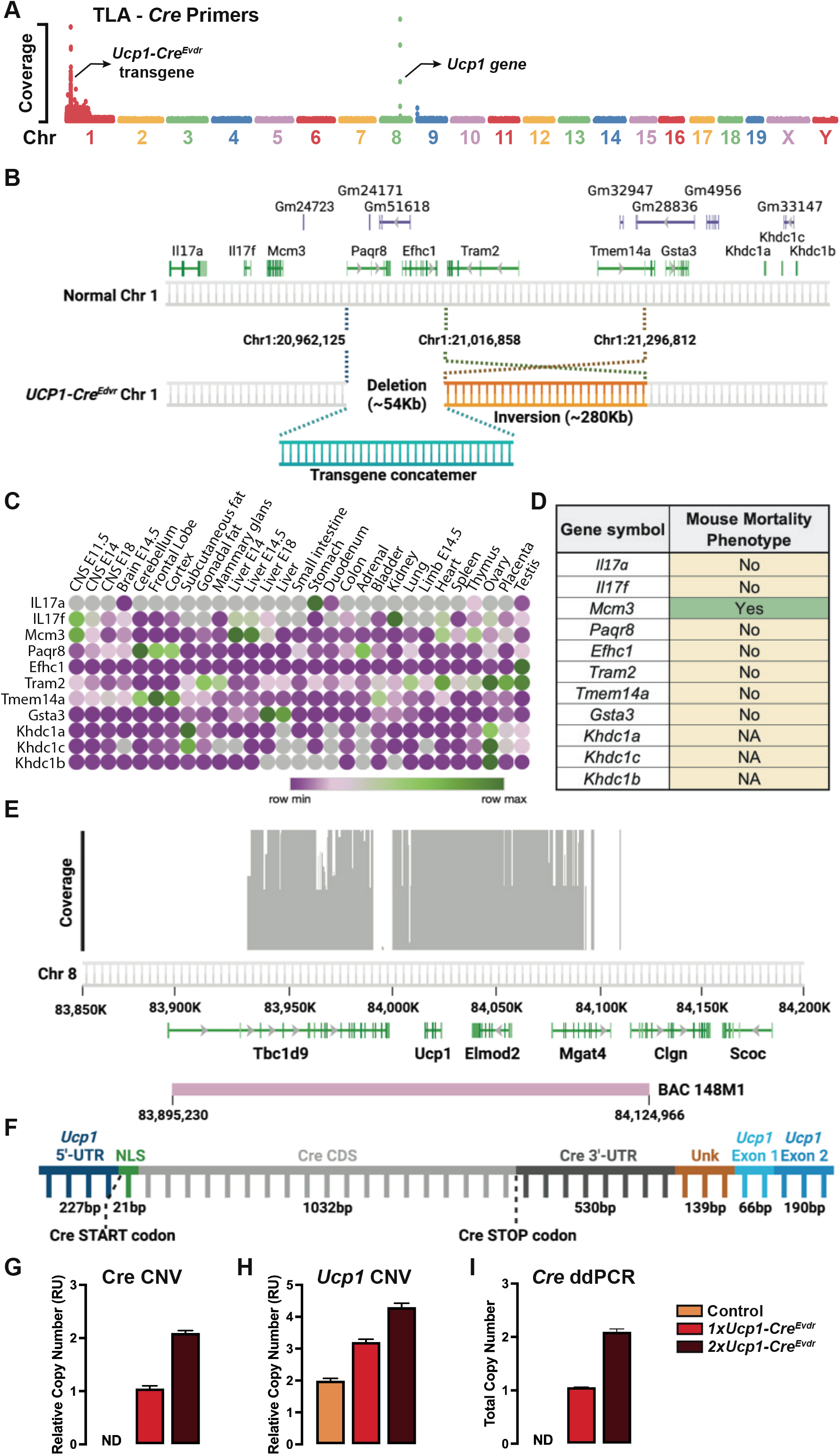
(A) Whole genome TLA mapping analysis of 1x*Ucp1-Cre^Evdr^* genome using primers specifics for the sequences of *Cre*. (B) Schematic representation of the identified integration site of the *Ucp1-Cre^Evdr^* transgene. (C) Gene expression of coding genes surrounding the *Ucp1-Cre^Evdr^* transgene from the Mouse ENCODE transcriptome data. (D) Knockout mortality phenotype association to each coding gene surrounding the *Ucp1-Cre^Evdr^* transgene in MGI. (E) Coverage of BAC 148M1 inserted within the *Ucp1-Cre^Evdr^* transgene as determined by TLA. (F) *de novo* reconstruction of the CRE-proximal part of *Ucp1-Cre^Evdr^* transgene mRNA from iBAT RNA-seq data. UTR: untranslated region; NLS: nuclear location signal; CDS: coding sequence; Unk: unknown. (G) Copy number assay of *Cre* of control, 1x*Ucp1-Cre^Evdr^* and 2x*Ucp1-Cre^Evdr^* mice. n= 3 per genotype. (H) Copy number assay of *Ucp1* of control, 1x*Ucp1-Cre^Evdr^* and 2x*Ucp1-Cre^Evdr^* mice. n= 8 per genotype. (I) Absolute copy number by ddPCR of *Cre* of control, 1x*Ucp1-Cre^Evdr^* and 2x*Ucp1-Cre^Evdr^* mice. n= 3 per genotype.

Next, we explored the *Ucp1-Cre^Evdr^*concatemer structure. To do this, we first analyzed the TLA data. We find that ∼75% of the original BAC used to generate the *Ucp1-Cre^Evdr^* mice (BAC 148M1), which covers ∼230Mb of chromosome 8 surrounding the *Ucp1* gene, is incorporated with the transgene in chromosome 1 [Figure 2E]. Notably, this includes an entire extra copy of the *Ucp1* gene. To further elucidate transgene structure, we performed *de novo* assembly of *Cre*-proximal transgene sequence of iBAT RNA-seq reads in *Ucp1-Cre^Evdr^* mice using the *Cre-recombinase* coding sequence as bait. Upstream of the Cre coding sequence, we identified the proximal *Ucp1* 5’UTR sequence followed by the start codon and SV40 nuclear localization signal [Figure 2F]. The *Cre* coding sequence is followed at 3’ by a 3’UTR and a short sequence of unknown function [Figure 2F]. The length of sequencing fragments limits the extend of the transgenic *Ucp1* gene we can detect as part of the *Ucp1-Cre^Evdr^* transgene unambiguously against the endogenous copy; however, we find *Cre* transgene bound to *Ucp1* exons 1 and part of exon 2 [Figure 2F]. The presence of *Ucp1* mRNA within the *Ucp1-Cre^Evdr^* transgene transcript suggests that the extra copy of *Ucp1* gene may be expressed.

TLA cannot discern the number of repetitions occurring within the concatemer. To clearly determine the copy number of *Ucp1* and *Cre* genes within the *Ucp1-Cre^Evdr^*concatemer, we employed two quantitative PCR-based techniques. First, we developed copy number assays against the *Ucp1* intron 3 to assess the number of copies of *Ucp1* gene in genomic DNA of controls, 1x*Ucp1-Cre^Evdr^* and 2x*Ucp1-Cre^Evdr^*mice. Control littermates were used as reference for two copies of *Ucp1* gene [Figure 2G-H]. In contrast, 1x*Ucp1-Cre^Evdr^* mice have three copies and 2x*Ucp1-Cre^Evdr^*mice have four copies of the *Ucp1* gene [Figure 2G-H]. However, this assay requires calibration with reference samples, limiting its ability to discern *Cre* copy number within the transgene concatemer. To solve this, we used digital droplet PCR (ddPCR)^39^ to quantify the absolute copy number of *Cre* in *HaeIII*-digested genomic DNA. As anticipated, control mice contained no copies of *Cre*. In contrast, 1x*Ucp1-Cre^Evdr^* mice harbored a single *Cre* copy, while 2x*Ucp1-Cre^Evdr^* mice contained two copies per genome [Figure 2I]. Taken together, these complementary assays indicate that the *Ucp1-Cre^Evdr^* transgene structure comprises one additional copy of the *Ucp1* gene and a single copy of *Cre*.

### *UCP1-Cre^Evdr^* Transgene Induces Profound Effects in BAT and WAT Biology

Because *Ucp1-Cre^Evdr^*directly affects fat size, we next examined if the *Ucp1-Cre^Evdr^*transgene directly impacts fat biology. First, we performed unbiased whole genome expression profiling of iBAT and psWAT from control, 1x*Ucp1-Cre^Evdr^*and 2x*Ucp1-Cre^Evdr^* female mice. Strikingly, the presence of the transgene profoundly alters the transcriptomic landscape of both fat depots. In 1x*Ucp1-Cre^Evdr^* iBAT, 1012 genes are upregulated and 905 downregulated compared to controls [Figure 3A]. Even more dramatic effects are evident in 1x*Ucp1-Cre^Evdr^* psWAT, with 3742 genes upregulated and 3130 downregulated despite barely detectable transgene expression [Figure 3B, S1N]. Comparisons between 2x*Ucp1-Cre^Evdr^* and control females reveal similar transcriptomic perturbations, with over 10-fold more altered genes in psWAT (8313) than in iBAT (760) [Figure 3C-D]. A lower number of genes are significantly different in iBAT and psWAT when comparing 1x*Ucp1-Cre^Evdr^*and 2x*Ucp1-Cre^Evdr^* [Figure S3A-B]. Remarkably, this suggest that the major effect appears from having just one copy of the *Ucp1-Cre^Evdr^* transgene. The dramatic transcriptomic effects in *Ucp1-Cre^Evdr^* fat depots, especially the psWAT, suggest either potent inter-tissue consequences or major effects from transgene insertion. In summary, unbiased transcriptional profiling indicates that the *Ucp1-Cre^Evdr^*transgene may profoundly impact the molecular state of both brown and white adipose tissues.

**Figure 3:**
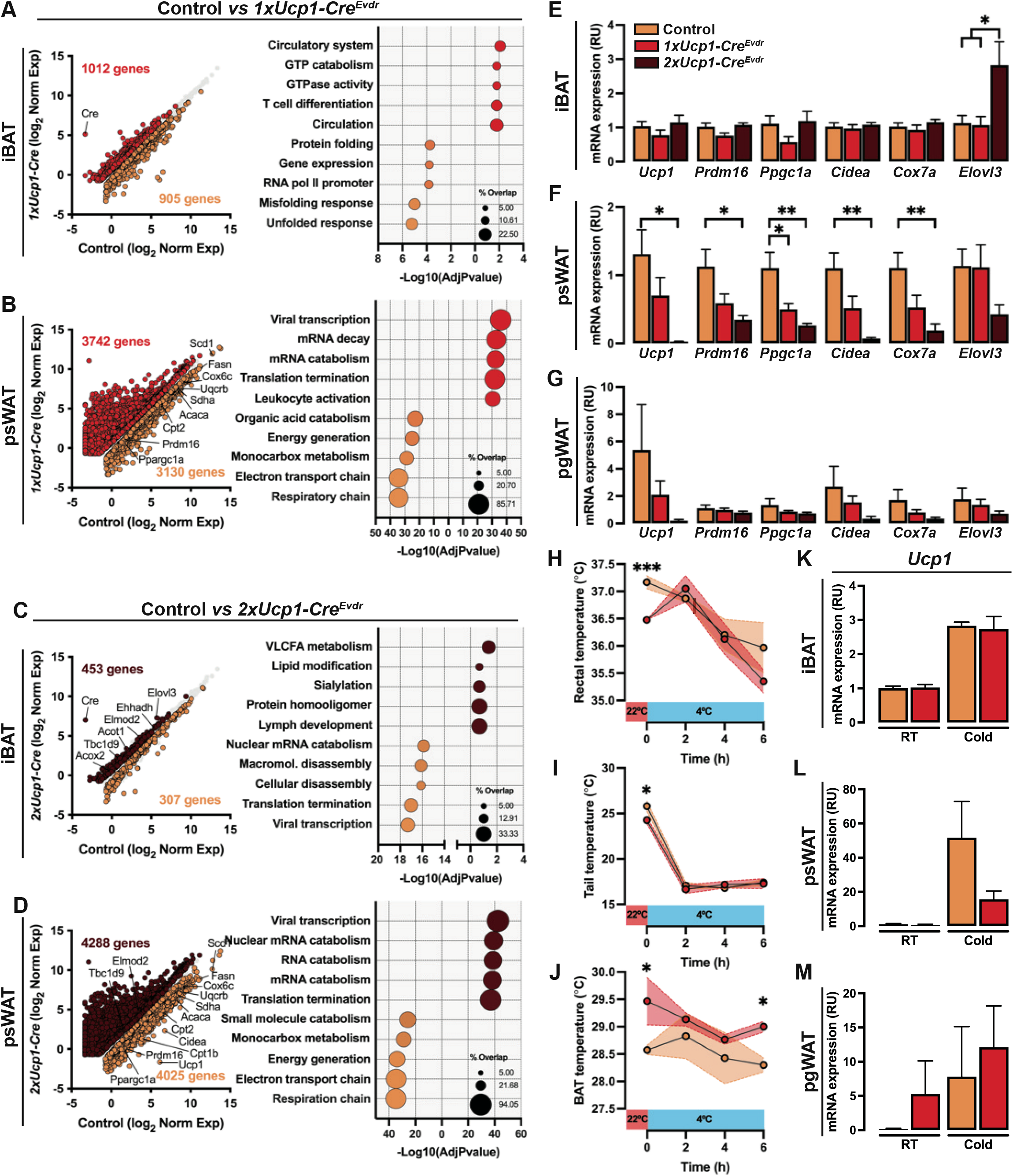
(A) RNA-seq comparing female control and 1x*Ucp1-Cre^Evdr^* iBAT gene expression (left). Each dot represents one gene. Corresponding GO analysis (right). Genes and pathways significantly enriched in controls are labeled in orange and those enriched in 1x*Ucp1-Cre^Evdr^* are labeled in red. (B) RNA-seq comparing female control and 1x*Ucp1-Cre^Evdr^* psWAT gene expression (left). Each dot represents one gene. Corresponding GO analysis (right). Genes and pathways significantly enriched in controls are labeled in orange and those enriched in 1x*Ucp1-Cre^Evdr^* are labeled in red. (C) RNA-seq comparing female control and 2x*Ucp1-Cre^Evdr^* iBAT gene expression (left). Each dot represents one gene. Corresponding GO analysis (right). Genes and pathways significantly enriched in controls are labeled in orange and those enriched in 2x*Ucp1-Cre^Evdr^* are labeled in brown. (D) RNA-seq comparing female control and 2x*Ucp1-Cre^Evdr^* psWAT gene expression (left). Each dot represents one gene. Corresponding GO analysis (right). Genes and pathways significantly enriched in controls are labeled in orange and those enriched in 2x*Ucp1-Cre^Evdr^* are labeled in brown. (E) qPCR analysis of iBAT of control, 1x*Ucp1-Cre^Evdr^* and 2x*Ucp1-Cre^Evdr^* females at 6 weeks of age. n= 6. (F) qPCR analysis of psWAT of control, 1x*Ucp1-Cre^Evdr^* and 2x*Ucp1-Cre^Evdr^* females at 6 weeks of age. n= 6. (G) qPCR analysis of pgWAT of control, 1x*Ucp1-Cre^Evdr^* and 2x*Ucp1-Cre^Evdr^* females at 6 weeks of age. n= 6. (H) Rectal temperature of control and 1x*Ucp1-Cre^Evdr^* females undergoing acute cold challenge. n= 3 controls, 4 1x*Ucp1-Cre^Evdr^*. Statistical significance was calculated using unpaired t-test within timepoint. (I) Tail temperature of control and 1x*Ucp1-Cre^Evdr^* females undergoing acute cold challenge. n= 3 controls, 4 1x*Ucp1-Cre^Evdr^*. Statistical significance was calculated using unpaired t-test within timepoint. (J) BAT temperature of control and 1x*Ucp1-Cre^Evdr^* females undergoing acute cold challenge. n= 3 controls, 4 1x*Ucp1-Cre^Evdr^*. Statistical significance was calculated using unpaired t-test within timepoint. (K) qPCR analysis of iBAT of control and 1x*Ucp1-Cre^Evdr^* females after cold challenge or maintained at room temperature. n= 3. Statistical significance was calculated using unpaired t-test between room temperature (RT) and cold samples. (L) qPCR analysis of psWAT of control and 1x*Ucp1-Cre^Evdr^* females after cold challenge or maintained at room temperature. n= 3. Statistical significance was calculated using unpaired t-test between room temperature (RT) and cold samples. (M) qPCR analysis of pgWAT of control and 1x*Ucp1-Cre^Evdr^* females after cold challenge or maintained at room temperature. n= 3. Statistical significance was calculated using unpaired t-test between room temperature (RT) and cold samples. Unless otherwise noted, data are mean + SEM and statistical significance was calculated using one-way ANOVA followed by Tukey’s multiple comparisons test. *P < 0.05, **P < 0.01, ***P < 0.001. For RNA-seq, differential genes were selected by false discovery rate (FDR) < 0.05 with no fold-change cut-off.

Pathway analysis of the significantly altered genes reveals downregulation of mitochondrial activity pathways (e.g., electron transport chain, respiratory chain, energy generation) in 1x*Ucp1-Cre^Evdr^* psWAT [Figure 3B]. Conversely, mRNA biology pathways are upregulated in 1x*Ucp1-Cre^Evdr^*psWAT [Figure 3B]. 1x*Ucp1-Cre^Evdr^* iBAT display irregularities in diverse pathways not directly related to energy metabolism [Figure 3A]. Similar patterns are observed in 2x*Ucp1-Cre^Evdr^* mice, with psWAT exhibiting suppressed energy generation pathways and elevated mRNA biology [Figure 3D]. In contrast to 1x*Ucp1-Cre^Evdr^* iBAT, 2x*Ucp1-Cre^Evdr^*iBAT uniquely show upregulation of lipid metabolism including very long chain fatty acid metabolism [Figure 3C]. iBAT of 2x*Ucp1-Cre^Evdr^*appears to be enriched in genes related to multiple terms of fatty acid metabolism while psWAT of 2x*Ucp1-Cre^Evdr^* is depleted of them [Figure S3A-B]. Collectively, these results poised a scenario in which a single copy of the *Ucp1-Cre^Evdr^*transgene may affect energy metabolism and thermogenesis pathways in iBAT and psWAT and these effects are heightened in 2x*Ucp1-Cre^Evdr^* mice.

We next used qPCR to verify whether thermogenic gene expression is affected by the *Ucp1-Cre^Evdr^*transgene. In iBAT, key markers of the classic thermogenesis pathway are largely unchanged across control, 1x*Ucp1-Cre^Evdr^* and 2x*Ucp1-Cre^Evdr^*females, apart from increased *Elovl3* in 2x*Ucp1-Cre^Evdr^*[Figure 3E]. In contrast, psWAT of 1x*Ucp1-Cre^Evdr^* females display an approximately 50% reduction in thermogenic genes (i.e., *Ucp1*, *Prdm16*, *Ppgc1a*, *Cidea*, *Cox7a*), but not *Elovl3* [Figure 3F]. These suppressions are amplified in 2x*Ucp1-Cre^Evdr^* psWAT with reductions of 98% in *Ucp1*, 69% in *Prdm16* and 94% in *Cidea* [Figure 3F]. Similar thermogenic depletion is evident in 2x*Ucp1-Cre^Evdr^* pgWAT of females, including 96% lower *Ucp1* expression on average [Figure 3G]. Male iBAT and psWAT, but not pgWAT, show similar results to females suggesting a gender specific effect on pgWAT [Figure S3C-E]. Collectively, these results indicate that the presence of the *Ucp1-Cre^Evdr^* transgene impairs expression of thermogenic genes in psWAT.

Given the profound effects of *Ucp1-Cre^Evdr^* on thermogenic gene expression, we hypothesized that the transgene alone, without additional genetic manipulation, would impact cold response. To test this, room temperature-acclimated control and 1x*Ucp1-Cre^Evdr^* female mice were exposed to 4°C for 6 hours. Intriguingly, 1x*Ucp1-Cre^Evdr^* mice exhibit elevated core body temperature, measured by a rectal thermometer, before cold exposure which normalized to control levels during cold [Figure 3H]. Under room temperature conditions, 1x*Ucp1-Cre^Evdr^* mice display lower tail but higher iBAT surface temperatures compared to controls [Figure 3I-J]. This suggests a scenario in which 1x*Ucp1-Cre^Evdr^* shows tail vasoconstriction and elevated iBAT thermogenesis as a physiological mechanism to increase body temperature. Upon cold exposure, 1x*Ucp1-Cre^Evdr^*tail temperature normalized while iBAT remain hyperactive [Figure 3I-J]. However, this is insufficient to maintain core temperature. Together, these data reveal dysfunctional thermogenic regulation and body temperature control in mice harboring just one copy of the *Ucp1-Cre^Evdr^*transgene.

We next investigated the effects of acute cold exposure on control and 1x*Ucp1-Cre^Evdr^* mice. After 6 hours of cold exposure, 1x*Ucp1-Cre^Evdr^* mice exhibit slightly greater body weight loss compared to controls [Figure S3F]. iBAT weight is unchanged between genotypes; however, psWAT and pgWAT are smaller in 1x*Ucp1-Cre^Evdr^*mice [Figure S3F], indicating increased lipid utilization to maintain body temperature. Cold-induced thermogenic gene expression is largely similar between control and 1x*Ucp1-Cre^Evdr^* iBAT, with comparable upregulation of *Ucp1* (∼2.8-fold), and *Elovl3* (∼4.2-fold) [Figure 3K, S3H]. However, psWAT displayed blunted activation, with *Ucp1* increasing ∼52-fold in controls but only ∼16-fold in 1x*Ucp1-Cre^Evdr^* by cold together with reduced upregulation of other markers such as *Elovl3* [Figure 3L, S3I]. pgWAT shows no differences after cold treatment between controls and 1x*Ucp1-Cre^Evdr^*mice [Figure 3M, S3J]. Histological analysis aligns with the gene expression data, revealing fewer multilocular adipocytes in 1x*Ucp1-Cre^Evdr^* psWAT after cold exposure, while iBAT and pgWAT appeared unaffected [Figure S3G]. Together, these data suggest an scenario of impaired psWAT thermogenic activation in response to acute cold stress in mice harboring the *Ucp1-Cre^Evdr^*transgene.

### *UCP1-Cre^Evdr^* transgene has the potential to express high levels of UCP1

The discovery of an additional transgenic *Ucp1* gene within the *Ucp1-Cre^Evdr^* transgene raised the question of its potential functionality. We hypothesized that this transgene derived *Ucp1* could contribute to overall UCP1 levels. However, the transgenic *Ucp1* sequence is identical to endogenous C57Bl6/J *Ucp1*, precluding its discrimination from the native genes. To overcome this limitation, we adopted a tissue-specific approach utilizing *Ucp1-floxed* mice (Ucp1tm1a(EUCOMM)Hmgu, EUCOMM)^40^ harboring LoxP sites flanking exon 2 [Figure S4A-B]. By crossing *Ucp1-floxed* mice with *Ucp1-Cre^Evdr^*animals, we would selectively ablate endogenous *Ucp1* while preserving the transgenic variant. This would allow us to assess the functional impact of the transgenic *Ucp1* gene within the *Ucp1-Cre^Evdr^*concatemer.

However, standard endpoint PCR genotyping failed to identify any mice homozygous for *Ucp1-floxed* (*Ucp1-fl/fl*) and positive for *Ucp1-Cre^Evdr^* (aka. *Ucp1-fl/fl^Ucp^*^1^*^-CreEvdr^* mice)[Fig 4A]. Concurrently, an excess proportion of heterozygous *Ucp1-floxed* (*Ucp1-fl/+*) carriers of *Ucp1-Cre^Evdr^* (aka. *Ucp1-fl/+^Ucp^*^1^*^-CreEvdr^* mice) was observed [Fig 4A]. As explained above, embryonic lethality due to *Ucp1* deficiency is not expected.

Thus, we ponder the hypothesis that the transgenic *Ucp1* gene within the *Ucp1-Cre^Evdr^*concatemer [Fig 2E] would mask the *floxed* status of the endogenous *Ucp1* alleles by yielding a wildtype band in endpoint PCR genotyping. To overcome this, we obtained the sequences surrounding the FRT site present in floxed allele (*tm1c*) of the Ucp1tm1a(EUCOMM)Hmgu mice [Figure S4A-B]. Next, we designed a copy number assay specific to detect this FRT sequence [Figure S4A-B]. We used wildtype, *Ucp1-fl/+* and *Ucp1-fl/fl* mice to calibrate our assay to zero, one and two copies of FRT [Figure 4B]. Using this approach, we readily distinguished *Ucp1-fl/+^Ucp^*^1^*^-CreEvdr^* mice harboring one FRT copy from *Ucp1-fl/fl^Ucp^*^1^*^-CreEvdr^*mice with two FRT copies [Figure 4B].

**Figure 4:**
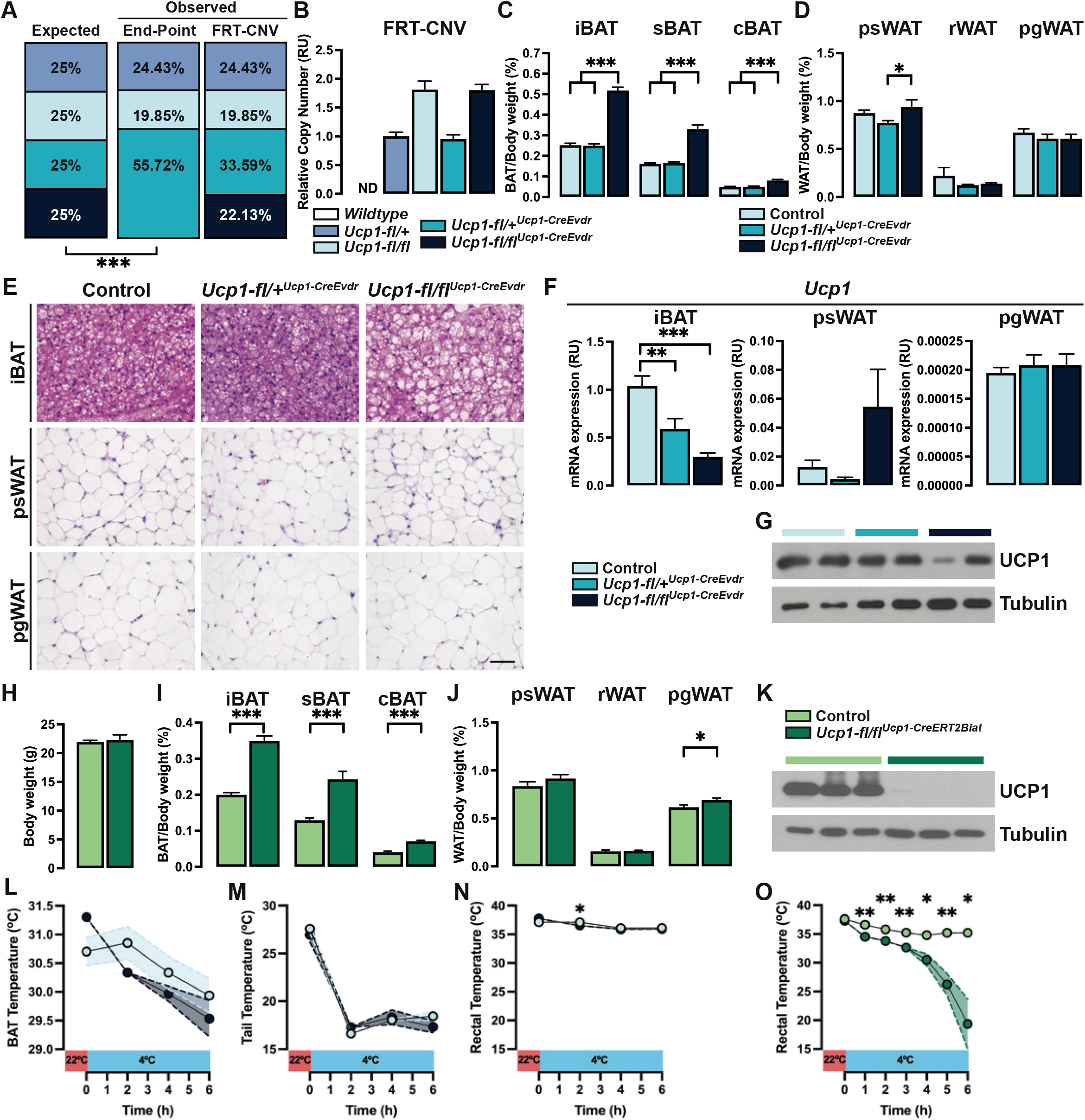
(A) Expected and observed offspring genotypes obtained from end-point PCR genotyping and *FRT* copy number assay. n=131 pups. Statistical significance was calculated using Chi-square test. (B) Copy number assay of *FRT*. n= 4. (C) BAT weights of control, *Ucp1-fl/+^Ucp^*^1^*^-CreEvdr^* and *Ucp1-fl/fl^Ucp^*^1^*^-CreEvdr^* males. n= 14 control, 13 *Ucp1-fl/+^Ucp^*^1^*^-CreEvdr^*, 9 *Ucp1-fl/fl^Ucp^*^1^*^-CreEvdr^*. (D) WAT weights of control, *Ucp1-fl/+^Ucp^*^1^*^-CreEvdr^* and *Ucp1-fl/fl^Ucp^*^1^*^-CreEvdr^* males. n= 14 control, 13 *Ucp1-fl/+^Ucp^*^1^*^-CreEvdr^*, 9 *Ucp1-fl/fl^Ucp^*^1^*^-CreEvdr^*. (E) Representative H&E images of fat depots from control, *Ucp1-fl/+^Ucp^*^1^*^-CreEvdr^* and *Ucp1-fl/fl^Ucp^*^1^*^-CreEvdr^* males. n= 4. Scale bar, 50μm. (F) qPCR analysis of *Ucp1* in adipose tissue depots of control, *Ucp1-fl/+^Ucp^*^1^*^-CreEvdr^* and *Ucp1-fl/fl^Ucp^*^1^*^-CreEvdr^* males. Values are relative to those of iBAT control. n= 8. (G)Western blot of iBAT protein lysates of control, *Ucp1-fl/+^Ucp^*^1^*^-CreEvdr^* and *Ucp1-_fl/fl_Ucp1-CreEvdr* _males._ (H) Body of control and *Ucp1-fl/fl^Ucp^*^1^*^-CreERT2Biat^* males. n= 7 control, 6 *Ucp1-fl/fl^Ucp^*^1^*^-^CreERT2Biat*. (I) BAT weights of control and *Ucp1-fl/fl^Ucp^*^1^*^-CreERT2Biat^* males. n= 7 control, 6 *Ucp1-_fl/fl_Ucp1-CreERT2Biat*. (J) WAT weights of control and *Ucp1-fl/fl^Ucp^*^1^*^-CreERT2Biat^* males. n= 7 control, 6 *Ucp1-_fl/fl_Ucp1-CreERT2Biat*. (K) Western blot of iBAT protein lysates of control and *Ucp1-fl/fl^Ucp^*^1^*^-CreERT2Biat^* males. (L) BAT temperature of control and *Ucp1-fl/fl^Ucp^*^1^*^-CreEvdr^* males undergoing acute cold challenge. n= 6 controls, 3 *Ucp1-fl/fl^Ucp^*^1^*^-CreEvdr^*. Statistical significance was calculated using unpaired t-test within timepoint. (M) Tail temperature of control and *Ucp1-fl/fl^Ucp^*^1^*^-CreEvdr^* males undergoing acute cold challenge. n= 6 controls, 3 *Ucp1-fl/fl^Ucp^*^1^*^-CreEvdr^*. Statistical significance was calculated using unpaired t-test within timepoint. (N) Rectal temperature of control and *Ucp1-fl/fl^Ucp^*^1^*^-CreEvdr^* males undergoing acute cold challenge. n= 6 controls, 3 *Ucp1-fl/fl^Ucp^*^1^*^-CreEvdr^*. Statistical significance was calculated using unpaired t-test within timepoint. (O) Rectal temperature of control and *Ucp1-fl/fl^Ucp^*^1^*^-CreERT2Biat^* males undergoing acute cold challenge. n= 3 controls, 3 *Ucp1-fl/fl^Ucp^*^1^*^-CreEvdr^*. Statistical significance was calculated using unpaired t-test within timepoint. Unless otherwise noted, data are mean + SEM. Statistical significance was calculated using unpaired t-test or one-way ANOVA followed by Tukey’s multiple comparisons test. *P < 0.05, **P < 0.01, ***P < 0.001.

Genotyping with the FRT copy number assay revealed expected Mendelian ratios of control, *Ucp1-fl/+^Ucp^*^1^*^-CreEvdr^*, and *Ucp1-fl/fl^Ucp^*^1^*^-CreEvdr^* progeny [Figure 4A], proving that this strategy overcomes the confounding effects from the transgenic *Ucp1* sequence. At 6 weeks of age, total body weight is equivalent between control, *Ucp1-fl/+^Ucp^*^1^*^-^*^CreEvdr^, and *Ucp1-fl/fl^Ucp^*^1^*^-CreEvdr^* mice [Figure S4C]. BAT depots weights are similar between controls and *Ucp1-fl/+^Ucp^*^1^*^-CreEvdr^* [Figure 4C]. However, *Ucp1-fl/fl^Ucp^*^1^*^-CreEvdr^* mice display marked ∼2-fold increase in weight in all BAT [Figure 4C]. WAT, liver or muscle tissue weights are unchanged across genotypes [Figure 4D, S4D-F]. In summary, targeted BAT-specific *Ucp1* ablation elicit pronounced BAT growth without impacting body and WAT weight.

Histological examination reveals the iBAT hypertrophy in *Ucp1-fl/fl^Ucp^*^1^*^-CreEvdr^* mice is attributable to uniformly enlarged brown adipocytes engorged with excessive lipid [Figure 4E]. In contrast, WAT depots morphology is largely unaffected by genotype [Figure 4E]. This shows a lack of morphological change compensation in WAT by the targeted deletion of *Ucp1* in BAT.

qPCR analysis reveals a ∼50% and ∼70% reduction in iBAT *Ucp1* mRNA in *Ucp1-fl/+^Ucp^*^1^*^-CreEvdr^* and *Ucp1-fl/fl^Ucp^*^1^*^-CreEvdr^* mice, respectively, compared to controls [Figure 4F]. *Cre* mRNA is equal in iBAT of *Ucp1-fl/+^Ucp^*^1^*^-CreEvdr^* and *Ucp1-fl/fl^Ucp^*^1^*^-CreEvdr^* mice [Figure S4G]. iBAT also shows a compensatory increased expression of classic thermogenic key markers *Ppargc1a*, *Cox7a*, and *Elovl3* [Figure S4J]. In psWAT, *Ucp1* expression is unaltered; however *Ucp1-fl/fl^Ucp^*^1^*^-CreEvdr^* mice display elevated expression of *Cidea*, *Cox7a*, and *Elovl3* [Figure 4F, S4K]. pgWAT gene expression is largely unchanged [Figure 4F, S4L]. *Cre* mRNA expression is slightly upregulated in psWAT and pgWAT of *Ucp1-fl/fl^Ucp^*^1^*^-CreEvdr^* mice partially recapitulating a possible compensation in psWAT, but with levels still much lower than those found in iBAT [Figure S4H-I]. Thus, BAT targeted *Ucp1* ablation induces depots-specific effects, including a rather unique selective compensatory thermogenic activation in iBAT and psWAT.

The high residual *Ucp1* mRNA expression in iBAT of *Ucp1-fl/fl^Ucp^*^1^*^-CreEvdr^*mice was unexpected. This is because, first, *Ucp1* is restricted to mature brown adipocytes and second, because *Ucp1-Cre^Evdr^* drives recombination in essentially all brown adipocytes, as we and others have previously shown^28,41^. To further investigate this, we examined UCP1 protein levels by western blot. Remarkably, iBAT lysates of *Ucp1-fl/fl^Ucp^*^1^*^-CreEvdr^* mice retained variable but high UCP1 protein levels, averaging approximately 70% of control [Figure 4G]. Given the broad *UCP1-Cre^Evdr^*-mediated excision in brown adipocytes, the substantial UCP1 retained seems unlikely to be derived from endogenous *Ucp1* genes. This paradoxical preservation of UCP1 protein suggests functionally significant expression from the transgenic *Ucp1* within the *UCP1-Cre^Evdr^* concatemer.

As a control, we crossed *Ucp1-floxed* mice with the tamoxifen-inducible *Ucp1-CreERT2^Biat^* allele. Tamoxifen treatment of *Ucp1-fl/fl^Ucp^*^1^*^-CreERT2Biat^* mice does not change body weight but increases BAT weights with a slight enlargement of pgWAT and liver, while other fats and tissues are unchanged [Figure 4H-J, S4M-O]. Critically, tamoxifen treatment leads to highly efficient ablation of UCP1 protein in iBAT [Figure 4K]. This confirms that the *Ucp1-floxed* allele can be efficiently deleted and reinforce the hypothesis that the lack of UCP1 deletion in *Ucp1-fl/fl^Ucp^*^1^*^-CreEvdr^* may arise from ectopic transgene expression.

To test for functionality, we performed acute cold challenges by exposing mice at 6°C for 6 hours. Intriguingly, *Ucp1-fl/fl^Ucp^*^1^*^-CreEvdr^*mice exhibit rather elevated core and BAT temperatures before starting cold exposure, compared to littermate controls [Figure 4L-N]. Unexpectedly, *Ucp1-fl/fl^Ucp^*^1^*^-CreEvdr^* mice are proficient in maintaining core body temperature during cold exposure at the same level than littermate controls [Figure 4N]. In stark contrast, *Ucp1-fl/fl^Ucp^*^1^*^-CreERT2Biat^* mice rapidly become hypothermic at 6°C, reflecting the efficient ablation of UCP1 in BAT [Figure 4O]. Together, these results indicate that the UCP1 protein present in *Ucp1-fl/fl^Ucp^*^1^*^-CreEvdr^* BAT, which may be bestowed by the *Ucp1-Cre^Evdr^* allele, may confer sufficient thermogenic capacity to preserve body temperature.

## Discussion

The Cre-Lox system is invaluable for spatial and temporal dissection gene function, tracing lineages, and labeling cells. However, the validation of *Cre-recombinase* transgenes in the literature is usually incomplete. In particular, the *UCP1-Cre^Evdr^*transgene transformed BAT and metabolism research. However, our findings reveal several key unexpected caveats of this widely used line including (1) increased mortality, growth defects, and craniofacial alterations in homozygosity; (2) substantial genomic disruptions; (3) profound impacts on BAT and psWAT function in both hemizygosity and homozygosity; and (4) potential misexpression of *Ucp1* itself under high thermogenic burden. These findings highlight the importance of carefully considering the validation of transgenic lines before embarking on experimental studies.

BAC Cre-drivers are usually validated only by examining spatiotemporal recombination, fueling comprehensive databases that guide selection among a large and growing number of *Cre-recombinases* available^4,5^. Causes for unexpected transgene expression include insertion effects on local regulation or integration of sequences leading to ectopic expression. Alternatively, unanticipated activity may reflect previously unknown endogenous gene expression. Adipocyte-targeting *Cre* lines exemplify these issues. The promoter of the fatty acid binding protein 4 gene, also known as adipocyte protein 2 (aP2), was used to generate two independent *aP2-Cre* mouse models with the intention to specifically target mature adipocytes^42,43^. However, these *aP2-Cre* models were found to inefficiently target mature adipocytes while exhibiting broad recombination in the brain, endothelial cell in adipose tissues, macrophages, adipocyte progenitors, and elsewhere^5,16,20,44–47^. This prompted development of *Cre* lines driven by the promoter of adiponectin^19,20^. However, *Ucp1-Cre^Evdr^*also exhibits widespread brain activity^41,48^, including regions controlling feeding and non-shivering thermogenesis^49–51^. Importantly, very low endogenous *Ucp1* expression partially overlaps with these brain areas^41,48^, suggesting *Ucp1-Cre^Evdr^* partially recaptures native *Ucp1* regulation. However, if this reflects endogenous *Ucp1* expression or expression of the *Ucp1* gene found within the *UCP1-Cre^Evdr^* transgene, that we find here, is not known at this point.

The recent close examination of some BAC transgenic lines, beyond cellular expression, has shown that the transgenes themselves can result in phenotypes leading to potential misinterpretations of the intended genetic modifications^52–54^. Random BAC transgene insertion is frequently associated with substantial genomic alterations, often disrupting gene coding sequences and creating small or large rearrangements^6,7,8^. As expected, *Ucp1-Cre^Evdr^* mice exhibit major structural variations at the insertion site including relatively large deletions and inversions. Historically, the position in the genome and the genomic alterations induced by the random insertion of a transgene have not been systematically examined. This is partially due to previous low-resolution techniques like FISH or linkage mapping which poorly defined insertion sites and structures. However, new sequencing approaches such as whole genome sequencing and TLA enable fine mapping of insertion locus, disruption effects, and integrated sequences^37^. For instance, whole genome sequencing and TLA both revealed *Adipoq-Cre^Evdr^* transgene inserted into the *Tbx18* gene on chromosome 9^19,55,56^, perturbing *Tbx18* expression and adding passenger gene copies with possible widespread effects^56^. Additionally, BAC transgenes normally integrate as concatemers leading to multiple full or partial copies of the transgene^3,57,58^. Using ddPCR, we find that *Cre* coding sequence is present in a single copy in *Ucp1-Cre^Evdr^* mice. This was also the case for the *Adipoq-Cre^Evdr^*transgene^56^. However, ddPCR analysis is limited to a small specific sequence via specific primers; thus, if partial coding or non-coding sequences are present, they may not be detected by ddPCR^39^. TLA is also not capable of defining the order or number of copies within the inserted concatemer in a transgenic line.

Defining insertion sites and the full structure including inserted sequences may only be possible by new long-range genome sequencing^7,59,60^. Additionally, transgenic strains are bred for generations, allowing accrual of modifications in transgenic sequences and the genetically linked endogenous sequences over time. Furthermore, *Cre*-drivers with initially robust expression can become leaky or lose activity. Monitoring integrated sequences over mouse line generations could enhance integrity of lines maintained and avoid misinterpretations.

The genetic alterations induced by random transgene insertion and the passenger sequences inserted can lead to confounding genetic effects that are dependent on the transgene rather than the intended genetic alteration. In *Ucp1-Cre^Evdr^* mice, we observe drastic transcriptional dysregulation in iBAT and psWAT, suggesting major changes in tissue function. Moreover, interactions between transgene effects and specific genetic manipulations (e.g., deletions) are unpredictable. While linking fat transcriptomics to whole body physiology is challenging, these data indicate that *Ucp1-Cre^Evdr^* the transgene has the potential to profoundly perturb adipose function. In this sense, 1x*Ucp1-Cre^Evdr^*mice show altered body temperature dynamics and distinct cold reactions when compared to controls. Compared to *Adipoq-Cre^Evdr^* mice, which show minimal adipose tissues gene expression changes^56^, the *Ucp1-Cre^Evdr^* effects are considerable. Beyond targeted tissues, transgenes can reprogram untargeted tissues due to genomic disruption and passenger genes. For *Ucp1-Cre^Evdr^*, the extra *Ucp1* gene seems highly expressed under high thermogenic demand, exemplifying how passenger genes can have unexpected impact. Overall, transgene insertion effects are diverse and context dependent. Thus, thorough characterization of each line is essential to parse transgene-specific artifacts from intended genetic effects.

Homozygosity often reveals phenotypes undetectable in heterozygotes, as most loss-of-function mutations display recessive inheritance. Thus, generating homozygous BAC transgenic models can uncover cryptic transgene-dependent effects. For example, crosses of hemizygous *Adipoq-Cre^Evdr^* mice have not produced homozygous mice suggesting lethality, although this experiment may have been underpowered^55^. The unexpected high mortality and other major effects caused by homozygosity of *Ucp1-Cre^Evdr^*imply impacts beyond adipose tissue. The multiple genes that are directly affected by the insertion site and that are expressed in a range of tissues may be responsible for these phenotypes. Moving forward, evaluating homozygous transgenic models, despite logistic challenges, may be a strong paradigm to ensure detection of subtle artifacts.

A limitation across published studies finding unexpected transgenic effects is that they usually lack mechanistic resolution. For example, the underlying causes of 2x*Ucp1-Cre^Evdr^* mortality and 1x*Ucp1-Cre^Evdr^* fat transcriptional changes remain unresolved. However, this knowledge gap in the literature is reasonable given the manifold possibilities. Effects could arise from chromosomal rearrangements, long-distance interactions, 3D conformation changes, or passenger sequences within the BAC, among other potential mechanisms. Moreover, unraveling any single mechanism may provide limited core insights into normal and pathogenic biology, as insertion effects likely stem from complex interactions only arising in the context of the transgenic line analyzed. While mechanistic details are invaluable, delineating the precise causes underlying insertion artifacts would require substantial efforts unlikely to significantly produce advancements. As such, the mechanistic ambiguity in these studies is expected.

New techniques for generating transgenic mice can mitigate issues with random BAC insertion, bolster rigor and drive discovery. CRISPR-directed insertion at known safe harbor *loci* provides control over transgene placement. Additionally, rational design of regulatory sequences enables precise spatiotemporal expression, avoiding complications from passenger DNA in large BAC constructs. As these and other innovations become widespread, they will complement and enhance previous data obtained with BAC transgenics.

The effects we report here may justify the reevaluation of some prior work using *Ucp1-Cre^Evdr^* mice to clarify confounding outcomes. However, such assessments will be complicated by the unpredictable interactions between the *Ucp1-Cre^Evdr^*effects and intended genetic changes. The strength and specificity of *Ucp1-Cre^Evdr^*transgene in brown adipocytes ensures that this line remains useful in verified contexts. In future studies, control groups with only the *Ucp1-Cre^Evdr^*transgene could be incorporated to parse its specific effects. Orthogonal tools like additional Cre lines (e.g., inducible *Ucp1-CreERT2^Biat^*allele) can also confirm results. Finally, new *Cre-recombinase* drivers may be generated, using methodologies described above, to confirm previous results. Though challenging, careful experimental design and layered validation can distinguish between effects dependent on the transgene, gene manipulation, or the interaction between the two.

In conclusion, validation of research tools is a requirement of several funding agencies, yet standards for doing so remain opaque. While BAC transgenics have revolutionized basic and biomedical research, limitations have become increasingly apparent. Overall, transparent validation, cautious interpretation, and technological innovations will maximize scientific rigor of BAC transgenics as future tools to catalyze discovery.

## Authors Contributions

Conceptualization and study design: JSG. Data collection: MH performed most experiments; OI, RM, JSG contribute to several experiments; CLBP performed skeletal staining and head measurements; MD performed ddPCR. Data analysis: MH, JSG. RNASeq data analysis: HWL. Data interpretation: JSG. Supervision: JSG; SAB supervised CLBP. Manuscript writing – Original draft: JSG. Manuscript writing – Review and editing: all authors. All authors approved the final manuscript.

## Acknowledgements

This work was supported by grants from the American Heart Association (18CDA34080527 to J.S.-G; and 19POST34380545 to RM), the NIH (R21OD031907 to J.S.-G, R35 DE027557 to SAB), a CCHMC Trustee Award to J.S.-G, a CCHMC Trustee Award to HL, a Center for Pediatric Genomics grant (CCHMC) to J.S.-G and HL and a Center for Mendelian Genetics grant (CCHMC) to J.S.-G. OI is supported by a Japanese Heart Foundation Research Abroad Award 2021. This project was supported in part by NIH P30 DK078392 of the Digestive Diseases Research Core Center in Cincinnati. This project was supported in part by the National Center for Advancing Translational Sciences of the National Institutes of Health, under Award Number 2UL1TR001425-05A1. The content is solely the responsibility of the authors and does not necessarily represent the official views of the NIH. We thank Helmholtz Zentrum München - Deutsches Forschungszentrum für Gesundheit und Umwelt (GmbH) for providing the mutant allele:C57BL/6N-Ucp1tm1a(EUCOMM)Hmgu, EMMA/INFRAFRONTIER (www.infrafrontier.eu) for distributing mouse line (EM:05767). We thank Christian Wolfrum for UCP1-CreERT2 mice. We thank Matt Kofron (Nikon Confocal Imaging Core at CCHMC) for support on image acquisition and analysis. We thank Barbara Cannon and Jan Nedergaard (The Wenner-Gren Institute, Stockholm University) for UCP1-floxed mice and for critically reading the manuscript. We thank Evan D Rosen (BIDMC, Harvard) for critically reading the manuscript. We thank all members of the Sanchez-Gurmaches lab for valuable discussions.

## STAR METHODS

### Lead contact and material availability

Further information and request for resources and reagents should be directed to and will be fulfilled by the Lead contact, Joan Sanchez-Gurmaches.

### Experimental model and subject details

#### Mice and mice housing

All mice used in this study were in C57Bl6/J background. UCP1-Cre (JAX stock 024670), R26R-mTmG mice (JAX stock 007676), Ucp1-CreER mice were described before^25^. Ucp1 flox mice were obtained from the EUCOMM program (C57BL/6N-Ucp1^tm1a^(EUCOMM)^Hmgu^/Ieg) after removal of the LacZ and neomycin cassette by Flippase.

Unless noted otherwise, mice were housed in the CCHMC Animal Medicine Facility in a clean room set at 22°C and 45% humidity on a daily 12h light/dark cycle, and kept in ventilated racks fed ad libitum with a standard chow diet, with bedding changed every two weeks. See figure legend for specific age and number of mice used. All animal experiments were approved by the CCHMC IACUC.

For long term temperature acclimation experiments, mice were housed in rodent incubators (7001-33 series, Caron) in pairs within the facilities of animal medicine of CCHMC. Room temperature group mice were co-housed in the same facility as the mice in rodent incubators. Mouse cages were changed weekly using components pre-adjusted to temperature. No cage enrichment was used in this set of experiments.

### Method details

#### MGI alleles and publications search

MGI database was searched with search word “transgenic” on 12/16/2022. Out of the 10,166 results, mice with Transgenic allele type were selected. Out of these, mouse models with location indicated as “unknown” were assigned as the location of the transgene unidentified group. The remainder mouse models were automatically assigned to transgene with known location group. To analyze Cre-driver mouse models the symbol of each mouse model was search for containing “Cre”. Out of the 1,968 mouse models found, location was assigned as above. Number of publications assigned to specific transgenic mice were found in MGI on July 2023.

#### GWAS associated traits and mouse mortality phenotype search

Mouse mortality phenotype associated to gene knockouts was searched in the MGI database. GWAS associated traits to specific genes were search in the Phenotype-Genotype Integrator of the NCBI.

#### Growth and Survival

Body weight and survival of mice was followed starting at week 3 of age as tail snips were taken for genotyping. Body weights and survival was recorded weekly after that until 6 weeks of age.

#### Tamoxifen treatment

Tamoxifen was dissolved in corn oil/ethanol (9:1 vol/vol) at 2mg/mL by shaking at 4°C overnight. 6 week old mice were injected with 2 mg/day/mouse for 5 times in a period of seven days. A subgroup of mice was additionally injected for four days during the third week after first injection. Mice were sacrificed three weeks after first injection.

#### Tissue dissection

Tissues were carefully dissected to avoid surrounding tissue contamination. Adipose tissue notation used here was described previously^61^. Mice were dissected at early morning without fasting or any other alteration, unless noted in the figure legend.

#### Acute cold exposure

Mice were placed at 4°C early in the morning of the experiment in overnight pre-chilled caging with free access to pre-chilled water with or without food.

#### Temperature measurements

Internal temperature was recorded by using a rectal thermometer probe (RET-3, Braintree Scientific Inc.). BAT and tail temperatures were obtained using an infrared thermal camera (FLIR T530 24°) in lightly anesthetized mice and analyzed with FLIR tools.

#### Tissue histology

Tissue pieces were fixed in 10% formalin. Embedding, sectioning and Hematoxylin and Eosin (H&E) staining was done by the CCHMC Pathology Core facility.

#### Craniofacial morphometric analysis

Samples were incubated in 0.005% Alizarin Red S (Sigma-Aldrich, A5533) in 1% KOH for 24 hours at room temperature and cleared in 1% KOH for 72 hours. Once cleared, samples were incubated in Glycerol:KOH 1% (50:50) solution. For imaging and long-term storage, samples were kept in 100% glycerol. Stained skulls were imaged using a Leica M165 FC stereo microscope system for measurements. The condylobasal length and the interorbital constriction length were measured in ImageJ. The ratio between them was used as skull shape defining factor.

#### qPCR analysis

Tissues were homogenized in a FASTPREP-24 (MP Biomedicals) using Qiazol (Qiagen). Total RNA was isolated using RNeasy kit (Qiagen), retrotranscription was done using High Capacity cDNA reverse transcription kit (#4368813, Applied Biosystems) and analyzed in a QuanStudio 3 real-time PCR machine (Thermofisher). Primer sequences are shown in Table S1.

#### Western blot analysis

Tissues were homogenized in a FASTPREP-24 (MP Biomedicals) and lysed in RIPA buffer (150 mM NaCl, 50 mM Hepes at pH 7.4, 0.1% SDS, 1% Triton X-100, 2 mM EDTA, 0.5% Na-deoxycholate) containing a protease and phosphatase inhibitor cocktail. Protein lysates (typically 10mg per lane) were mixed with 5X SDS sample buffer and run in SDS acrylamide/bis-acrylamide gels (typically 10 or 12%), transferred to PVDF membranes and detected with specific antibodies as specified in Table S2.

#### Copy number assays

Genomic DNA was isolated from tails or liver using the DNeasy Blood and Tissue kit spin columns (Qiagen) and diluted to 1 ng/μL. Copy number assays were done using Taqman ® copy number assays (ThermoFisher) using predesigned oligonucleotides assays using *Tfrc* as reference gene (ThermoFisher)(Table S4). qPCR was performed on a QuantStudio 6Flex real-time PCR system using the following protocol: 95°C for 10min followed by 40 cycles of 95°C for 15 s and 60°C for 60 s with manual Ct threshold at 0.2 and Autobaseline on. Results were analyzed on Copy Caller 2.0 software (ThermoFisher).

#### Digital droplet PCR

Reaction mixture was composed of 10 uL 2x ddPCR Supermix (without dUTPs; Bio-Rad, Hercules CA), 1 uL each of the proves against housekeeping and target gene (Bio-Rad)(Table S4), 1uL of *HaeIII* (NEB, R0108), 50 ng of DNA template, and adjusted to a final volume of 20 uL. Droplets were generated in a 96-well polypropylene plate (Bio-Rad) using the QX200 droplet generator (Bio-Rad). The plate containing the water-in-oil emulsions was sealed with foil using a PX1 PCR Plate Sealer (Bio-Rad) and placed in a C1000 Touch Thermal Cycler (Bio-Rad). The following conditions were used for amplification: 95°C for 10 minutes, 94°C for 30 seconds and 60°C for 1 minute (40 cycles 2°C/sec ramp rate), a 10-minute hold at 98°C, and a final hold at 4°C. The plate was processed using the QX200 droplet reader (Bio-Rad). Results were analyzed using QuantaSoft Analysis Pro software version 1.0.596.

#### Targeted Locus Amplification

UCP1-Cre transgene location and genetic rearrangements associated were approximated by Targeted Locus Amplification (TLA)(Cergentis B.V.) in splenocytes. Splenocytes were isolated from 8 weeks old UCP1-Cre hemizygous mice by pushing the spleen through a 40 mm mesh and collecting in 10% fetal bovine serum in PBS. After red blood cell lysis and washes with 10% fetal bovine serum in PBS, around 10 million spleen cells were aliquoted in cryovials in freezing media (10% fetal bovine serum, 10% DMSO, in PBS). TLA analysis was performed by Cergentis B.V. as previously reported^37^ with six independent pairs of primers (Table S3) using mouse mm10 genome as host reference.

#### Whole genome gene expression profile

RNA-seq reads were aligned to UCSC mouse genome 10 mm using STAR aligner^62^. Only uniquely aligned reads were used for downstream analysis. Raw read counts for each gene were measured using FeatureCounts in the subread package^63^ with an option, “-s 2 -O --fracOverlap 0.8”. Differential gene expression analysis was performed using EdgeR^64^. Genes with Fold-change > 1.5 and FDR < 0.05 were selected as differentially expressed genes. Gene ontology analysis was performed using Enrichr^65^.

#### *De novo* assembly

To build a *Cre* transgene sequence, we performed incremental alignment and *de novo* assembly. Initially, we built a STAR reference combining mm10 and the known *Cre* CDS sequence. Read 1 and 2 from each RNA-seq sample were aligned to the combined reference separately, where we selected read pairs for *de novo* assembly if at least one out of the pair is aligned to the *Cre* reference. And then, we pooled the selected read pairs and performed de novo assembly using Trinity^66^. Given the design of *Cre* transgene, we anticipated that the assembled sequence should fully cover *Cre* CDS and span from *Cre* CDS sequence to the 5’ half and 3’half of the Ucp1 exon #1.

We observed that the very first assembly results fully cover the *Cre* CDS and connect between Ucp1 5’ half and *Cre* CDS with a small gap of unknown sequence but not the 3’half. Therefore, we updated the *Cre* reference with the assembled sequence and repeated these steps of alignment, selection, and de novo assembly until the assembly result reaches the 3’ half of the Ucp1 exon #1, which happened after 4th round. The final assembly results were assessed and annotated using known references and Blast^67^.

#### Figure design

Figures were made in Adobe Illustrator. Several figures were created with BioRender.com.

## QUANTIFICATION AND STATISTICAL ANALYSIS

Data are presented as mean+s.e.m., unless stated otherwise. Unpaired t-test, analysis of variance (one or two ways) followed by Tukey’s multiple comparisons, Chi-square and Log-rank (Mantel-Cox) as appropriate, were used to determine statistical significance. No pre-test was used to choose sample size. Statistical analysis was done using GraphPad Prism except for global RNA expression (see methods). The number of mice used per experiment is stated in each figure legend. In all panels, *P < 0.05, **P < 0.01, ***P < 0.001.

---

**Figure S1:**
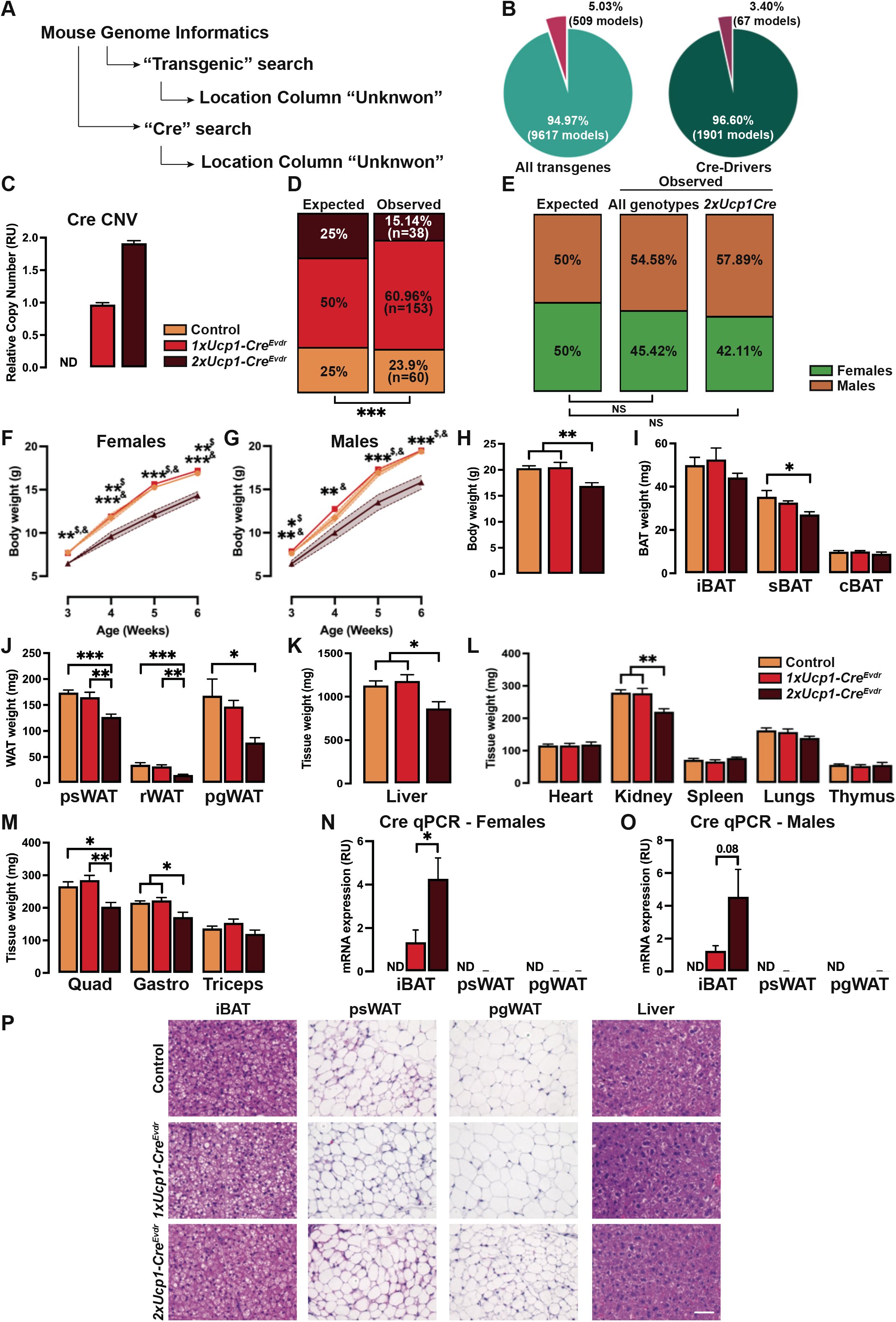
(A) Experimental strategy for the identification of known and unknown location for transgenes in Mouse Genome Informatics (MGI) site. (B) Proportions of all transgenes (left) and Cre-drivers (right) transgenic mice with known location in the genome. (C) Copy number assay of Cre of control, 1x*Ucp1-Cre^Evdr^* and 2x*Ucp1-Cre^Evdr^* mice. n= 5 per genotype. (D) Expected and observed offspring genotypes obtained from 1x*Ucp1-Cre^Evdr^* to 1x*Ucp1-Cre^Evdr^* crosses. N=251 pups from 46 litters. Statistical significance was calculated using Chi-square test. ***P < 0.001. (E) Expected and observed offspring sex distribution obtained from 1x*Ucp1-Cre^Evdr^* to 1x*Ucp1-Cre^Evdr^* crosses. N=251 pups from 46 litters. Statistical significance was calculated using Chi-square test. (F) Growth curves of control, 1x*Ucp1-Cre^Evdr^* and 2x*Ucp1-Cre^Evdr^* females. n= 17 controls, 41 1x*Ucp1-Cre^Evd^*, 6 2x*Ucp1-Cre^Evd^*. All 2x*Ucp1-Cre^Evd^* females analyzed in this growth curve survived up to week 6. $ indicates significant differences between 2x*Ucp1-Cre^Evdr^* and control, & indicates significant differences between 2x*Ucp1-Cre^Evdr^* and 1x*Ucp1-Cre^Evdr^*. (G) Growth curves of control, 1x*Ucp1-Cre^Evdr^* and 2x*Ucp1-Cre^Evdr^* males. n= 23 controls, 46 1x*Ucp1-Cre^Evdr^*, 14 2x*Ucp1-Cre^Evdr^*. All 2x*Ucp1-Cre^Evdr^* males analyzed in this growth curve survived up to week 6. $ indicates significant differences between 2x*Ucp1-Cre^Evdr^* and control, & indicates significant differences between 2x*Ucp1-Cre^Evdr^* and 1x*Ucp1-Cre^Evdr^*. (H) Body weights of control, 1x*Ucp1-Cre^Evdr^* and 2x*Ucp1-Cre^Evdr^* males. n= 6 controls, 6 1x*Ucp1-Cre^Evdr^*, 7 2x*Ucp1-Cre^Evdr^*. (I) BAT weights of control, 1x*Ucp1-Cre^Evdr^* and 2x*Ucp1-Cre^Evdr^* males. n= 6 controls, 6 1x*Ucp1-Cre^Evdr^*, 7 2x*Ucp1-Cre^Evdr^*. (J) WAT weights of control, 1x*Ucp1-Cre^Evdr^* and 2x*Ucp1-Cre^Evdr^* males. n= 6 controls, 6 1x*Ucp1-Cre^Evdr^*, 7 2x*Ucp1-Cre^Evdr^*. (K) Liver weight of control, 1x*Ucp1-Cre^Evdr^* and 2x*Ucp1-Cre^Evdr^* males. n= 6 controls, 6 1x*Ucp1-Cre^Evdr^*, 7 2x*Ucp1-Cre^Evdr^*. (L) Other organs weight of control, 1x*Ucp1-Cre^Evdr^* and 2x*Ucp1-Cre^Evdr^* males. n= 6 controls, 6 1x*Ucp1-Cre^Evdr^*, 7 2x*Ucp1-Cre^Evdr^*. (M) Skeletal muscles weight of control, 1x*Ucp1-Cre^Evdr^* and 2x*Ucp1-Cre^Evdr^* males. n= 6 controls, 6 1x*Ucp1-Cre^Evdr^*, 7 2x*Ucp1-Cre^Evdr^*. (N) qPCR analysis of *Cre* in adipose tissue depots of control, 1x*Ucp1-Cre^Evdr^* and 2x*Ucp1-Cre^Evdr^* females. n= 6. (O) qPCR analysis of *Cre* in adipose tissue depots of control, 1x*Ucp1-Cre^Evdr^* and 2x*Ucp1-Cre^Evdr^* males. n= 6. (P) Representative H&E images of fat depots and liver from control, 1x*Ucp1-Cre^Evdr^* and 2x*Ucp1-Cre^Evdr^* males. n= 4. Scale bar, 50μm. Unless otherwise noted, data are mean + SEM and statistical significance was calculated using one-way ANOVA followed by Tukey’s multiple comparisons test. *P < 0.05, **P < 0.01, ***P < 0.001.

**Figure S2:**
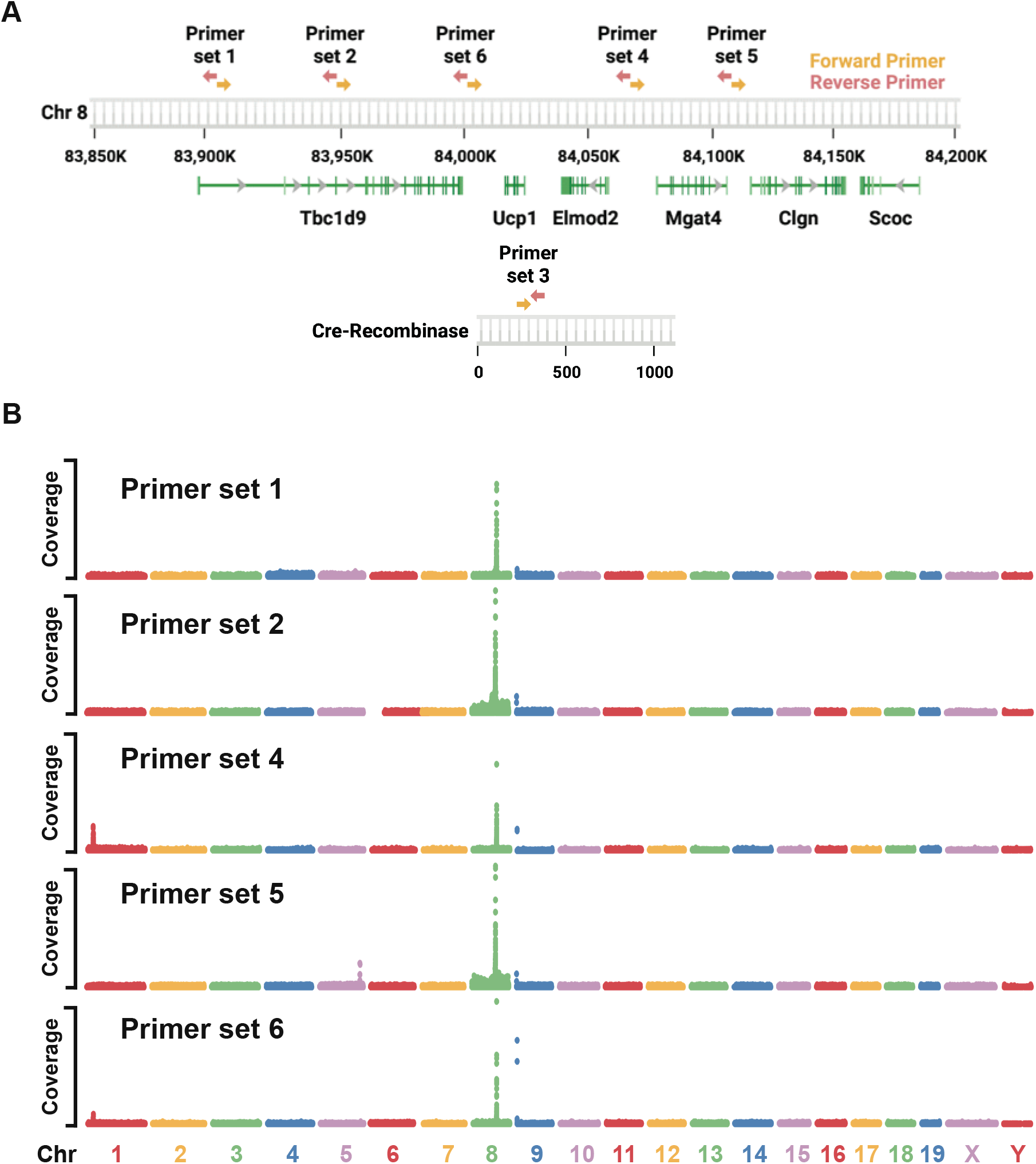
(A) Schematic representation of the location of the six probe pairs used for TLA analysis. See also Table S3. (B) Whole genome TLA mapping analysis of 1x*Ucp1-Cre^Evdr^* genome using probes surrounding the BAC 148M1 sequence.

**Figure S3:**
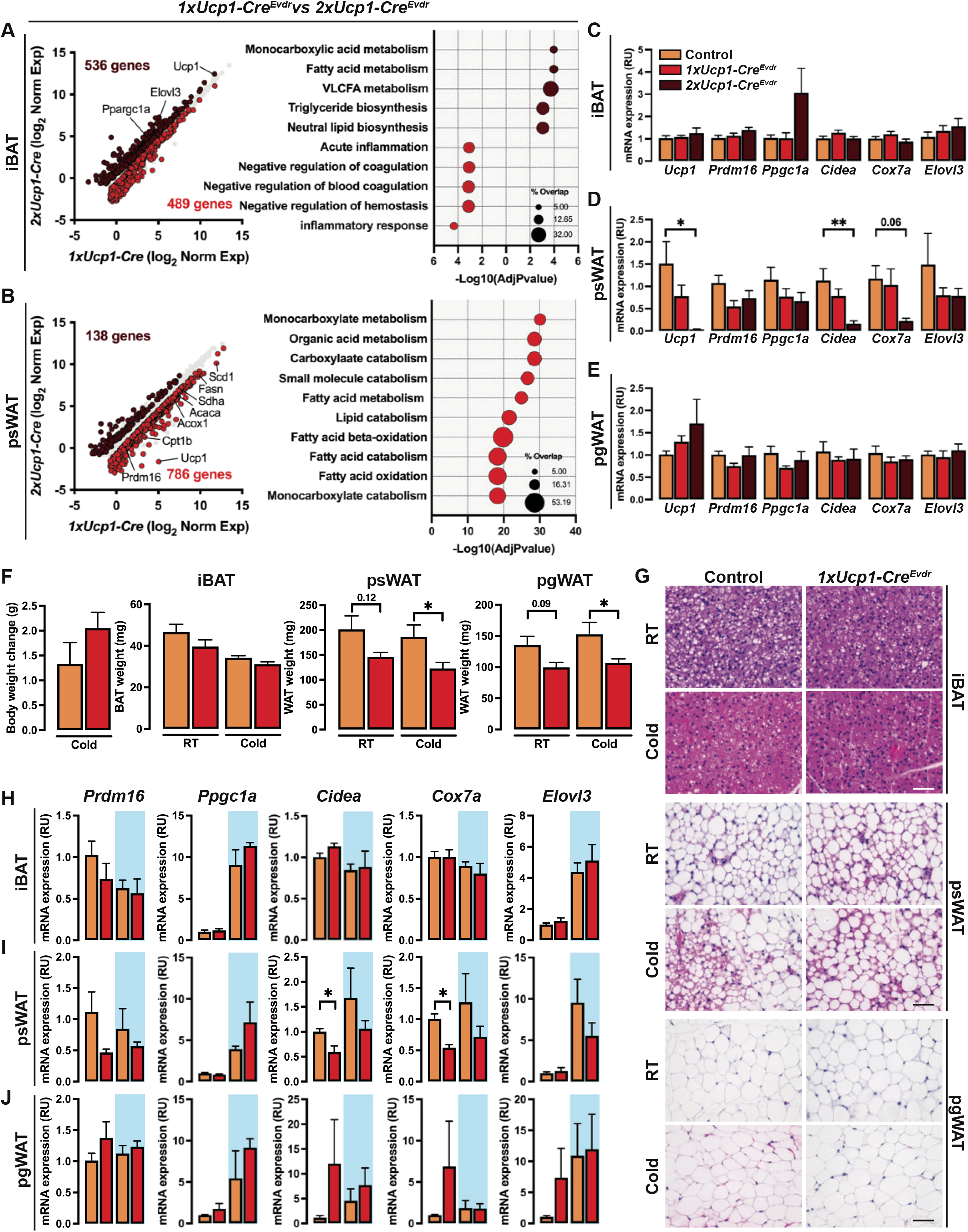
(A) RNA-seq comparing female 1x*Ucp1-Cre^Evdr^* and 2x*Ucp1-Cre^Evdr^* iBAT gene expression (left). Each dot represents one gene. Corresponding GO analysis (right). Genes and pathways significantly enriched in 1x*Ucp1-Cre^Evdr^* are labeled in red and those enriched in 2x*Ucp1-Cre^Evdr^* are labeled in brown. (B) RNA-seq comparing female 1x*Ucp1-Cre^Evdr^* and 2x*Ucp1-Cre^Evdr^* psWAT gene expression (left). Each dot represents one gene. Corresponding GO analysis (right). Genes and pathways significantly enriched in 1x*Ucp1-Cre^Evdr^* are labeled in red and those enriched in 2x*Ucp1-Cre^Evdr^* are labeled in brown. (C) qPCR analysis of iBAT of control, 1x*Ucp1-Cre^Evdr^* and 2x*Ucp1-Cre^Evdr^* males at 6 weeks of age. n= 6. (D) qPCR analysis of psWAT of control, 1x*Ucp1-Cre^Evdr^* and 2x*Ucp1-Cre^Evdr^* males at 6 weeks of age. n= 6. (E) qPCR analysis of pgWAT of control, 1x*Ucp1-Cre^Evdr^* and 2x*Ucp1-Cre^Evdr^* males at 6 weeks of age. n= 6. (F) Body weight change and tissue weights of control and 1x*Ucp1-Cre^Evdr^* females after cold challenge. n= 3 control RT, 3 1x*Ucp1-Cre^Evdr^* RT, 3 control cold, 4 1x*Ucp1-Cre^Evdr^* cold. Statistical significance was calculated using unpaired t-test between RT and cold samples. (G) Representative H&E images of fat depots from control, 1x*Ucp1-Cre^Evdr^* females after cold challenge. n= 4. Scale bar, 50μm. (H) qPCR analysis of iBAT of control and 1x*Ucp1-Cre^Evdr^* females after cold challenge or maintained at room temperature. n= 3. Statistical significance was calculated using unpaired t-test between RT and cold samples. (I) qPCR analysis of psWAT of control and 1x*Ucp1-Cre^Evdr^* females after cold challenge or maintained at room temperature. n= 3. Statistical significance was calculated using unpaired t-test between RT and cold samples. (J) qPCR analysis of pgWAT of control and 1x*Ucp1-Cre^Evdr^* females after cold challenge or maintained at room temperature. n= 3. Statistical significance was calculated using unpaired t-test between RT and cold samples. Unless otherwise noted, data are mean + SEM and statistical significance was calculated using one-way ANOVA followed by Tukey’s multiple comparisons test. *P < 0.05, **P < 0.01, ***P < 0.001.

**Figure S4:**
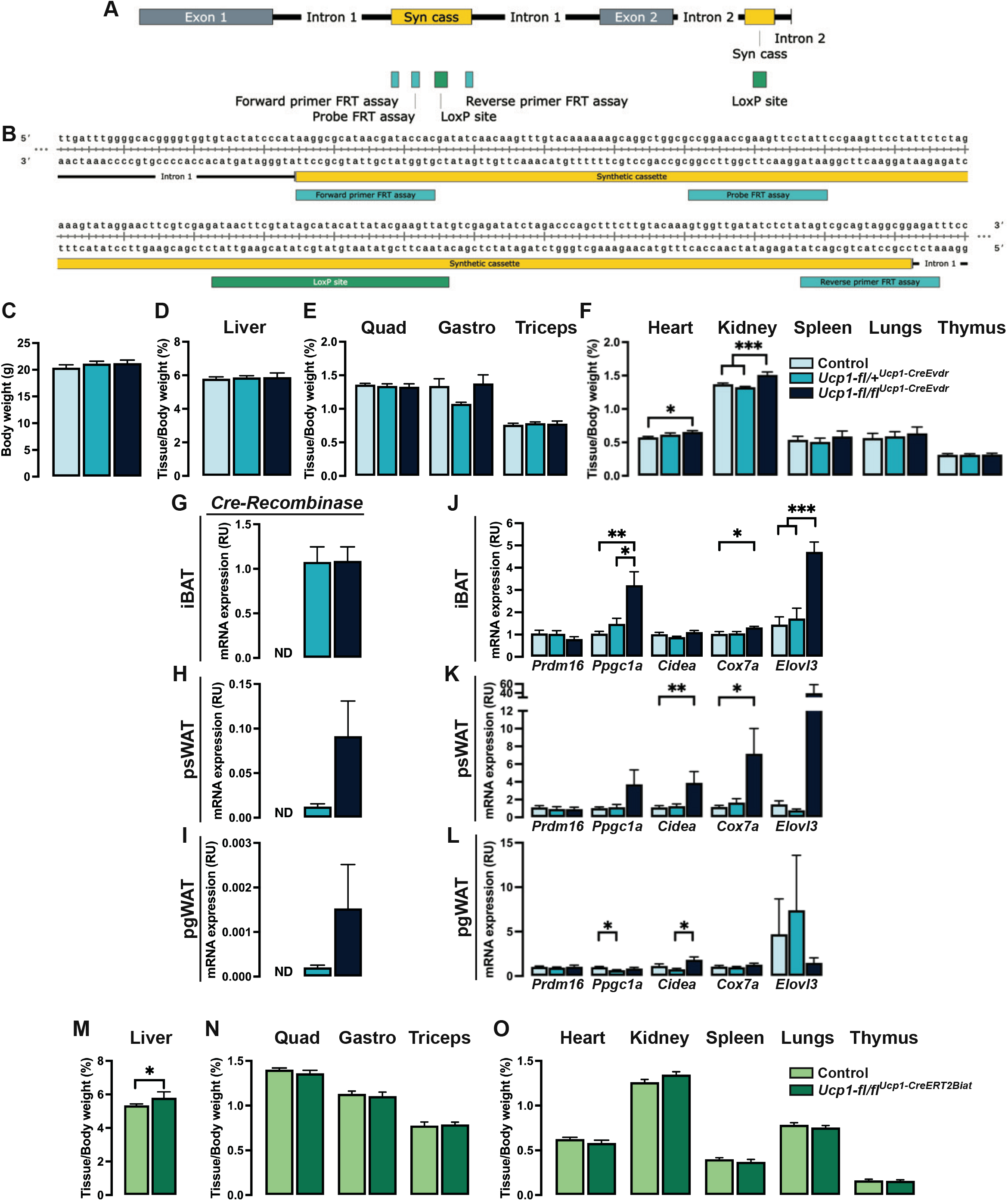
(A) Schematic representation of the genomic structure of the *Ucp1-floxed* (Ucp1tm1a(EUCOMM)Hmgu) allele. (B) Portion of the genomic sequence used for development of a specific FRT copy number assay. (C) Body weight of control, *Ucp1-fl/+^Ucp^*^1^*^-CreEvdr^* and *Ucp1-fl/fl^Ucp^*^1^*^-CreEvdr^* males. n= 14 control, 13 *Ucp1-fl/+^Ucp^*^1^*^-CreEvdr^*, 9 *Ucp1-fl/fl^Ucp^*^1^*^-CreEvdr^*. (D) Liver weight of control, *Ucp1-fl/+^Ucp^*^1^*^-CreEvdr^* and *Ucp1-fl/fl^Ucp^*^1^*^-CreEvdr^* males. n= 14 control, 13 *Ucp1-fl/+^Ucp^*^1^*^-CreEvdr^*, 9 *Ucp1-fl/fl^Ucp^*^1^*^-CreEvdr^*. (E) Muscle weights of control, *Ucp1-fl/+^Ucp^*^1^*^-CreEvdr^* and *Ucp1-fl/fl^Ucp^*^1^*^-CreEvdr^* males. n= 14 control, 13 *Ucp1-fl/+^Ucp^*^1^*^-CreEvdr^*, 9 *Ucp1-fl/fl^Ucp^*^1^*^-CreEvdr^*. (F) Other organs weight of control, *Ucp1-fl/+^Ucp^*^1^*^-CreEvdr^* and *Ucp1-fl/fl^Ucp^*^1^*^-CreEvdr^* males. n= 14 control, 13 *Ucp1-fl/+^Ucp^*^1^*^-CreEvdr^*, 9 *Ucp1-fl/fl^Ucp^*^1^*^-CreEvdr^*. (G) qPCR analysis of *Cre* in iBAT of control, *Ucp1-fl/+^Ucp^*^1^*^-CreEvdr^* and *Ucp1-fl/fl^Ucp^*^1^*^-CreEvdr^* males. n=8. (H) qPCR analysis of *Cre* in psWAT of control, *Ucp1-fl/+^Ucp^*^1^*^-CreEvdr^* and *Ucp1-fl/fl^Ucp^*^1^*^-CreEvdr^* males. Values are relative to those of iBAT control. n=8. (I) qPCR analysis of *Cre* in pgWAT of control, *Ucp1-fl/+^Ucp^*^1^*^-CreEvdr^* and *Ucp1-fl/fl^Ucp^*^1^*^-CreEvdr^* males. Values are relative to those of iBAT control. n=8. (J) qPCR analysis in iBAT of control, *Ucp1-fl/+^Ucp^*^1^*^-CreEvdr^* and *Ucp1-fl/fl^Ucp^*^1^*^-CreEvdr^* males. n=8. (K) qPCR analysis in psWAT of control, *Ucp1-fl/+^Ucp^*^1^*^-CreEvdr^* and *Ucp1-fl/fl^Ucp^*^1^*^-CreEvdr^* males. n=8. (L) qPCR analysis in pgWAT of control, *Ucp1-fl/+^Ucp^*^1^*^-CreEvdr^* and *Ucp1-fl/fl^Ucp^*^1^*^-CreEvdr^* males. n=8. (M) Liver weight of control and *Ucp1-fl/fl^Ucp^*^1^*^-CreERT2Biat^* males. n= 7 control, 6 *Ucp1-_fl/fl_Ucp1-CreERT2Biat*. (N) Muscles weight of control and *Ucp1-fl/fl^Ucp^*^1^*^-CreERT2Biat^* males. n= 7 control, 6 *Ucp1-_fl/fl_Ucp1-CreERT2Biat*. (O) Other organs weight of control and *Ucp1-fl/fl^Ucp^*^1^*^-CreERT2Biat^* males. n= 7 control, 6 *_Ucp1-fl/fl_Ucp1-reERT2Biat*. Unless otherwise noted, data are mean + SEM. Statistical significance was calculated using unpaired t-test or one-way ANOVA followed by Tukey’s multiple comparisons test. *P < 0.05, **P < 0.01, ***P < 0.001.

**Table S1.**
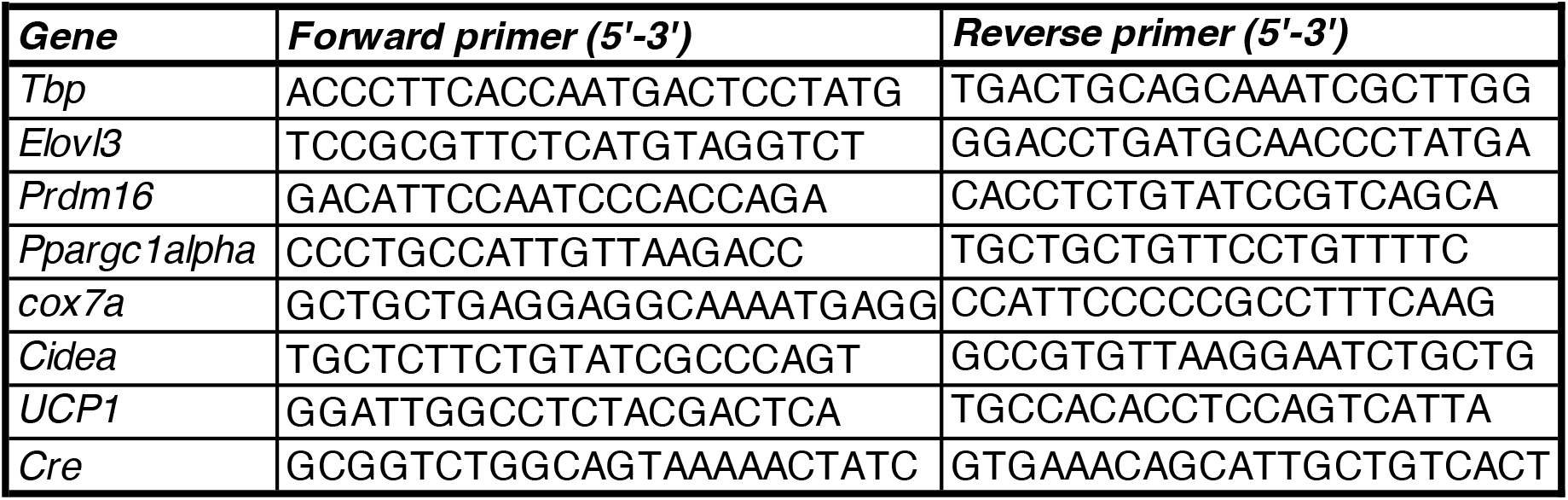
Mouse primers.

**Table S2.** Antibodies.

| Table S2. Antibodies |  |  |
| --- | --- | --- |
| **Antigen** | **Source** | **Concentration** |
| *UCP1* | Abcam (ab10983) | 1:1000 |
| *Tubulin* | Cell Signaling (cs2125) | 1:1000 |

**Table S3.**
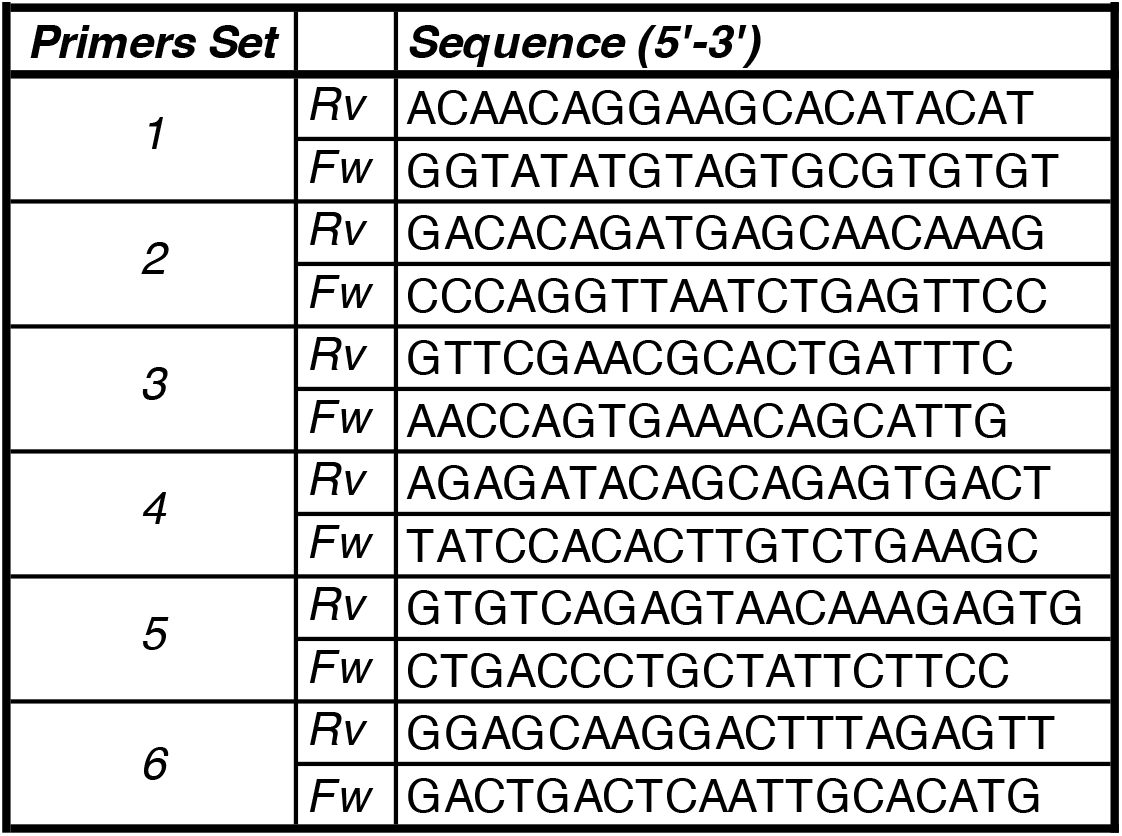
TLA Primer Sets.

**Table S4.**
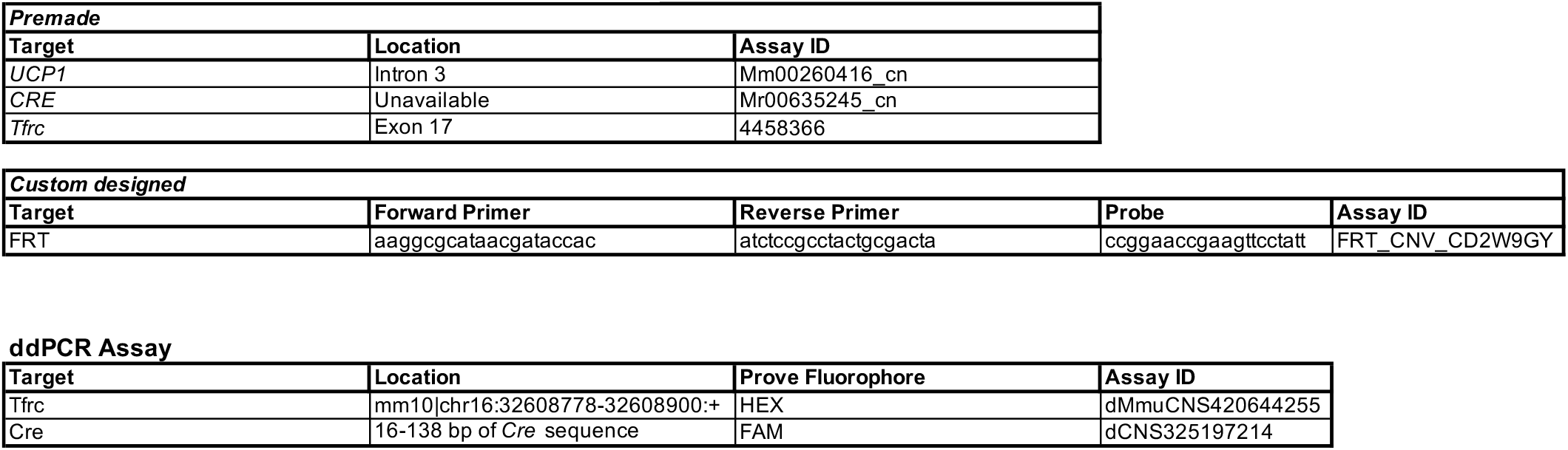
Copy Number Assays.

